# PD-1 checkpoint blockade disrupts CD4 T cell regulated adaptive B cell tolerance to foreign antigens

**DOI:** 10.1101/2021.06.10.447979

**Authors:** Chad R. Dufaud, Andrew G. Shuparski, Brett W. Higgins, Louise J. McHeyzer-Williams, Michael G. McHeyzer-Williams

## Abstract

Adaptive B cell immunity to environmental antigens must be regulated by multiple CD4 T cell dependent tolerance mechanisms. Using integrated single cell strategies, we demonstrate that acute PD-1 blockade induces extensive and selective local anti-inflammatory IgG1 plasma cell (PC) differentiation. Expansion of pre-existing IgG1 germinal center (GC) B cell and enhanced GC programming without memory B cell involvement reveals an isotype-specific GC checkpoint that blocks steady-state IgG1 antibody maturation. While there was no adjuvant impact on immunization, acute PD-1 checkpoint blockade exaggerates anti-commensal IgG1 antibody production, alters microbiome composition and exerts its action in a CD4 T cell dependent manner. These findings reveal a PD-1 controlled adaptive B cell tolerance checkpoint that selectively constrains maturation of pre-existing anti-inflammatory antibodies to prevent over-reaction to steady-state foreign antigens.

**In Brief:** PD-1 controls an adaptive B cell tolerance checkpoint in steady-state germinal centers to inhibit the maturation and production of IgG1 antibody with pre-existing foreign specificities.

**Highlights:**

– Acute PD-1 blockade induces extensive IgG1 PC differentiation at homeostasis
– PD-1 blockade releases an IgG1 GC B cell checkpoint that drives expansion and PC formation
– No adjuvant effect on foreign antigen but expansion of pre-existing IgG1 specificities to non-self
– PD-1 exerts CD4 T cell dependent tolerance in the GC to restrict IgG1 maturation to non-self

**Graphical Abstract:** 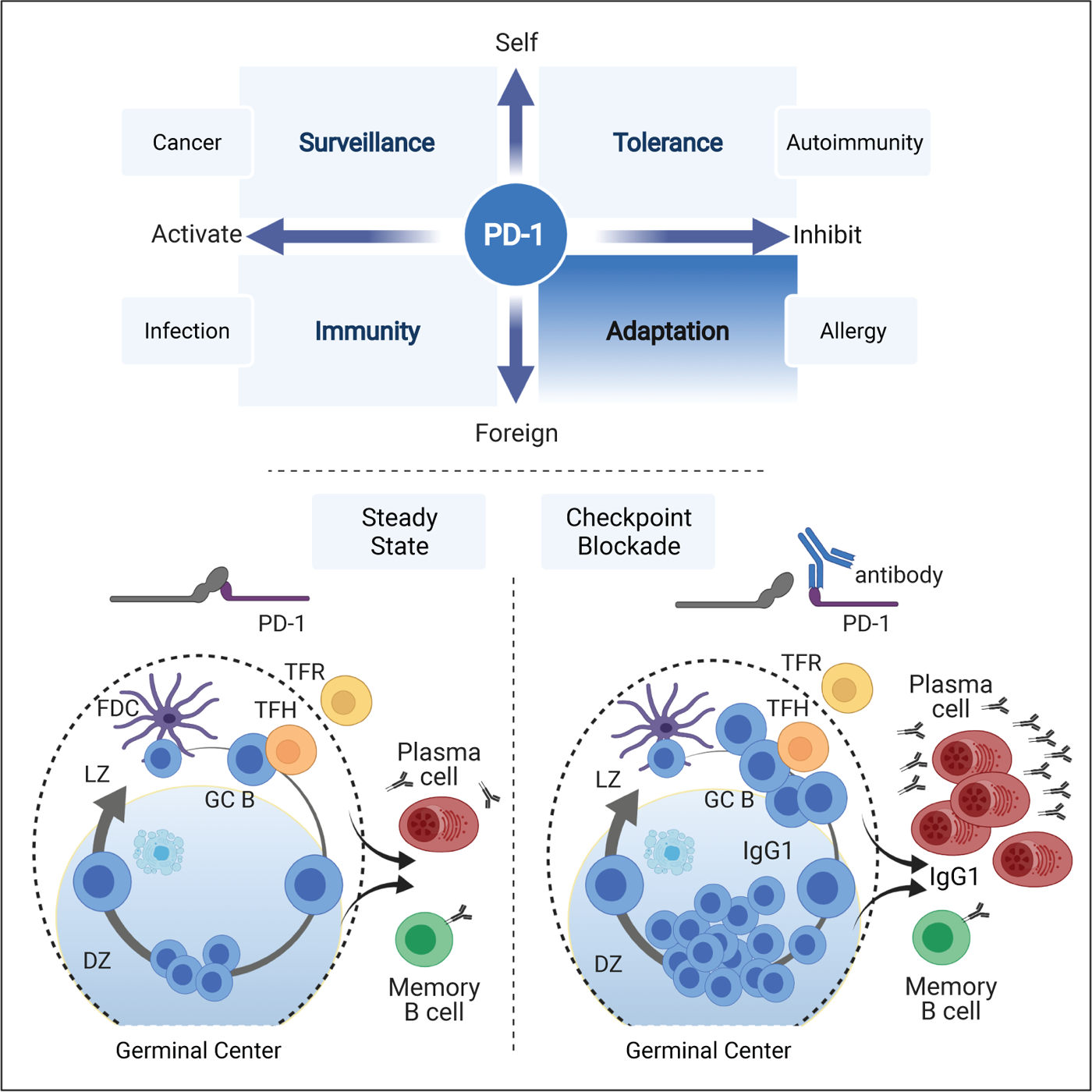

## INTRODUCTION

Adaptive B cell immunity involves a range of highly orchestrated cell fate decisions dynamically controlled by specialized CD4 follicular helper T (TFH) cell compartments (Crotty, 2019; Cyster and Allen, 2019). These TFH cells induce antibody class switch upon antigen-activation and promote affinity maturation within GC B cells to support long-term immune protection against a multitude of foreign antigens (McHeyzer-Williams et al., 2012; Mesin et al., 2016; Roco et al., 2019; Victora et al., 2010; Vinuesa et al., 2016). During development, some aspects of these CD4-regulated B cell compartments must also become tolerate to commensal organisms and a broad array of benign environmental antigens (Donaldson et al., 2018; Li et al., 2020; Morais et al., 2021; Yilmaz et al., 2021). Immune checkpoints regulate the quality and magnitude of CD4 T cell mediated signals that ultimately dictate B cell fate and impact immune function (ElTanbouly and Noelle, 2021; Liu et al., 2021; Qi, 2016; Sharpe and Pauken, 2018; Shi et al., 2018; Tan et al., 2021). It remains important to dissect how these immune checkpoints regulate adaptive B cell tolerance to environmental antigens.

PD-1 checkpoint blockade is a highly effective immune therapeutic that re-activates exhausted CD8 T cells during persistent infection (Barber et al., 2006; Hashimoto et al., 2018) and across a variety of cancers (Sharma et al., 2021; Wei et al., 2017). Despite their non-exhausted state, TFH cells (Ansel et al., 1999; Fazilleau et al., 2009; Vinuesa et al., 2005) and their follicular regulatory counterpart (TFR) (Gonzalez-Figueroa et al., 2021; Linterman et al., 2011; Wu et al., 2020) both express PD-1 with highest levels on GC localized subsets (Haynes et al., 2007; Shulman et al., 2013). Genetic ablation of PD-1 results in increased TFH and TFR cell expansion following immunization (Good-Jacobson et al., 2010; Hams et al., 2011; Sage et al., 2013), in the gut (Kawamoto et al., 2012) and in autoimmune models (Nishimura et al., 1999; Nishimura et al., 2001). These germline PD-1 deficiencies have variable impact on the magnitude, maturation and outcome of GC B cell selection (Good-Jacobson et al., 2010; Hams et al., 2011; Sage et al., 2016; Shi et al., 2018) with TFR-focused deletion influencing early pre-GC B cell activity and immune tolerance as well as reducing B cell recall responses (Clement et al., 2019; Lu et al., 2021; Tan et al., 2021).

Acute PD-1 blockade can enhance B cell responses (Dyavar Shetty et al., 2012; Mylvaganam et al., 2018; Velu et al., 2009) and promote TFH and B cell presence in the tumor microenvironment (Hollern et al., 2019; Sharonov et al., 2020) with a positive correlation to the outcome of immune therapy (Cabrita et al., 2020; Helmink et al., 2020; Petitprez et al., 2020). However, multiple adverse immune reactions to PD-1 blockade can involve autoimmune reactivity and inflammatory responses targeting barrier surfaces such as the gut (Dougan et al., 2021; Pauken et al., 2019). Identifying cellular pathways and steady-state molecular targets for PD-1 blockade remains an important fundamental need with high potential utility for understanding the mechanism of action as well as these adverse reactions of immune therapy.

Here, we use high-fidelity single cell sorting and targeted RNA sequencing to reveal the cellular and molecular mechanisms of acute PD-1 blockade on adaptive B cell tolerance. This CD4 T cell-dependent mechanism dysregulates pre-existing anti-inflammatory IgG1 GC B cell maturation to produce massive local expansion and differentiation of IgG1 PC and antibodies. Released specificities target pre-existing foreign antigens with no evidence for autoimmunity or overt adjuvant action in a vaccine context. We propose that this sub-class specific GC B cell checkpoint attenuates the maturation of pre-existing foreign specificities to avoid over-reacting to environmental antigens and maintain immune homeostasis.

## RESULTS

### Acute PD-1 blockade induces extensive IgG1 PC differentiation

To assess the role of the PD-1 pathway on CD4-regulated B cell immunity, we focus on IgG class-switched B cell fate and function in the steady-state spleen. Six days after single anti-PD-1 treatment across three different blocking antibody clones, we observed significant increases in IgG1 antibody-secreting cells in the spleen by functional analysis (**Figure 1A and 1B**). Direct assessment of isotype-specific PC numbers *in vivo* by high resolution flow cytometry revealed a rapid and selective increase in splenic IgG1-expressing PC (**Figure 1C - 1F**). Thus, without intentional immunization acute PD-1 blockade alone substantially disrupted isotype-specific antibody homeostasis and significantly enhanced the size of the local IgG1 PC compartment.

**Figure 1.**
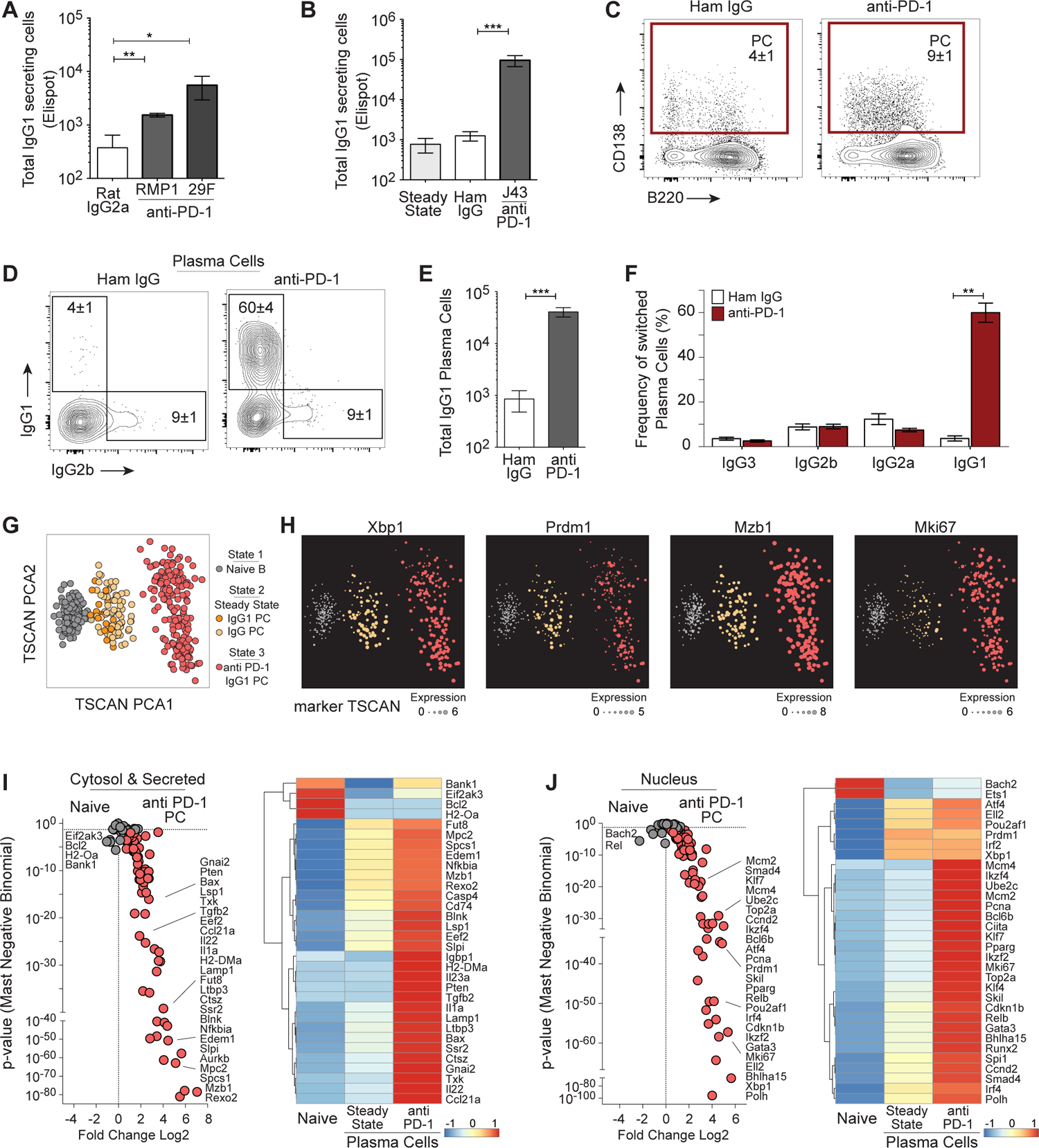
Acute PD-1 blockade induces extensive IgG1 Plasma Cell differentiation. **(A)** Total IgG1 secreting splenic cells by Elispot from mice treated 6 days prior with isotype control (Rat IgG2a, clone 2A3, n=4) or anti-PD-1 antibodies (clones, RMP1-14 or 29F.1A12, n=4 each), mean±SEM. **(B)** Untreated mice (Steady State, n=4) and mice treated 6 days prior with isotype control (Hamster IgG, n=4) or anti-PD-1 (clone J43, n=4) total IgG1 secreting cells, mean±SEM. **(C)** Flow cytometric analysis of class-switched Plasma Cells (PC: IgD-IgM-CD138+) and **(D)** distribution of IgG1 and IgG2b with **(E)** quantification for IgG1 (n=9) and **(F)** distribution of IgG sub-classes, (n=6) mean±SEM. **(G)** TSCAN pseudotime analysis of qtSEQ generated transcriptional programming from single FACS sorted Naïve B cells, IgG PC from Steady State, and IgG1 PC from anti-PD1 treated mice with **(H)** marker TSCAN gene expression. See also Figure S1. **(I)** Volcano plot of Naïve B cells compared to anti-PD1 IgG1 PC with averaged single cell heatmap (including Steady State PC) from cytosol and secreted gene products and **(J)** nucleus. See Figure S1 for cell membrane gene products and single cell heatmaps.

To discriminate between recent PC migration and rapid PC differentiation we sorted IgG-expressing splenic PC before and after checkpoint blockade for gene expression analysis. Individual cells from these rare compartments were uniquely bar-coded during cDNA synthesis, and gene targeted amplification of ∼500 mRNA species for sequencing utilizing digital deconvolution for single cell quantification (quantitative and targeted RNA sequencing; **qtSEQ**) (**Figure S1A - S1D**). By avoiding over-abundant immunoglobulin (Ig) mRNA, this single cell qtSEQ strategy provided clear access to many elements of the class-switched PC transcriptional program (**Figure S1E - S1G**). Pseudo-time TSCAN re-construction (Ji and Ji, 2016) orders and clearly separates naive B cells, steady-state IgG PC and anti-PD-1 treated IgG1 PC based on their targeted transcriptional profile (**Figure 1G**). Relative expression levels per cell of central PC-associated transcription factors (TF) *Xbp-1* and *Prdm-1* are similar between steady-state and anti-PD-1 treated PC, while proliferation associated *Mki67* and Ig co-chaperone *Mzb-1* are markedly up-regulated after treatment indicating recent cell expansion and IgG1 PC differentiation (**Figure 1H**).

Closer scrutiny of the induced IgG1 PC program reveals a multitude of transcriptional changes related to the steady-state IgG PC program that are significantly enhanced by acute checkpoint blockade. IgG1 PCs exhibit enhanced capacity for inter-cellular communication with increased expression of adhesion molecules (*Cd44, Cd48, Spn, Lgals1, Cd9*), SLAM family members (*Slamf6, Slamf9*), and TNF ligand and receptor superfamily members (*Tnfsf5, Tnfsf9, Tnfsf18, Tnfrsf4, Tnfrsf13c*) (**Figure S1H**). Acute changes in expression for antigen presentation (*H2.Ab1, Ly75, Cd81*), cell fate determination (*Notch2, Plxdn1*) and a range of known inhibitory receptors (*Havcr1, Timd2*) and co-stimulators (*Cd69, Cd28*, Cd24a) indicate major recent alterations in IgG1 PC development and function.

Enhanced expression of the intracellular machinery managing endoplasmic reticuluum (ER; *Mzb1, Spsc1, Ssr2, Edem1*), protein modification (*Slpi, Fut8*, *Txk*), chaperone function (*Igbp1*), signaling (*Blk, Nfkbia, Pten*), antigen processing (*H2.Dma, Cd74, Lamp1*) and metabolism (*Ctsz, Mpc2*) along with changes in secreted factors (*Il1a, Il22, Il23a, Ccl21a, Ltbp3*) were evident after treatment (**Figure 1I and S1I**). Highly significant increases in expression across multiple TF families (C2H2ZF: *Bcl6b, Ikzf2, Ikzf4, Klf4, Klf7;* WHforkhead: *Irf2, Irf4, Spib*) as well as upregulation of gene indicating recent PC differentiation activity (*Xbp1, Prdm1, Pou2af1, Bhlha15*) and PC response programs (*Gata3, Sta5b, Smad4, Skil*) were also induced by PD-1 checkpoint blockade (**Figure 1J and S1J**). Most informative are the dominant increases in cell cycle machinery (*Ccdn2, Cdkn1b, Pcna, Ube2c, Mki67*) and substantial elevation of DNA replication and repair elements (*Mcm2, Mcm4, Top2a, Polh*). Overall, these data indicate that PD-1 blockade drove recent and selective expansion of a PC precursor and then differentiation into a new anti-inflammatory IgG1 secreting PC compartment.

### IgG1 GC B cell expansion without memory B cell involvement

Class-switched IgG-expressing memory and GC B cells can be found in the steady-state spleen, but these small fractions of IgG1^+^ memory (GL7^-^CD38^+^) B cells remained unchanged after PD-1 blockade (**Figure 2A to 2C**). In contrast, there was a 30-50 fold selective expansion of IgG1-expressing GC (GL7^+^CD38^-^) B cells (**Figure 2D to 2F**). Significant IgG1 GC B cell or PC increases were not seen before day 6 after treatment (**Figure S2A and S2B**), with no evidence for early B cell activation (GL7^+^CD38^+^) or recruitment of IgG1^+^ memory B cells throughout treatment (data not shown). Single cell qtSEQ analysis with pseudo-time reconstruction (using TSCAN) clearly separates the transcriptional impact of checkpoint blockade from the steady-state IgG1 GC B cell program (**Figure S2C to S2E**; **Figure 2G**) with enhancement of major GC molecular program modifiers (*Bcl6, Aicda*) and evidence of recent proliferation (*Mki67*)(**Figure 2H**). There was a shift in GC B cell transcriptional activity with significant changes in expression for surface molecules involved in modifying BCR signals (*Cd79b, H2.Aa, Ptprc, Cd81*), TFH contact (*Cd244, Cd276, Spn, Havcr1*) and TNF receptor/ligand superfamily (*Tnfsf9, Tnfrsf25, Tnfrsf8*)(**Figure 2I and S2F**). There were also increases in expression of intracellular signaling components (*Dock8, Blnk, Aurkb, Blk*) and antigen processing machinery (*Cd74, H2.Dmb2*) that may accompany heightened selection activity of the GC cycle (**Figure S2G**).

**Figure 2.**
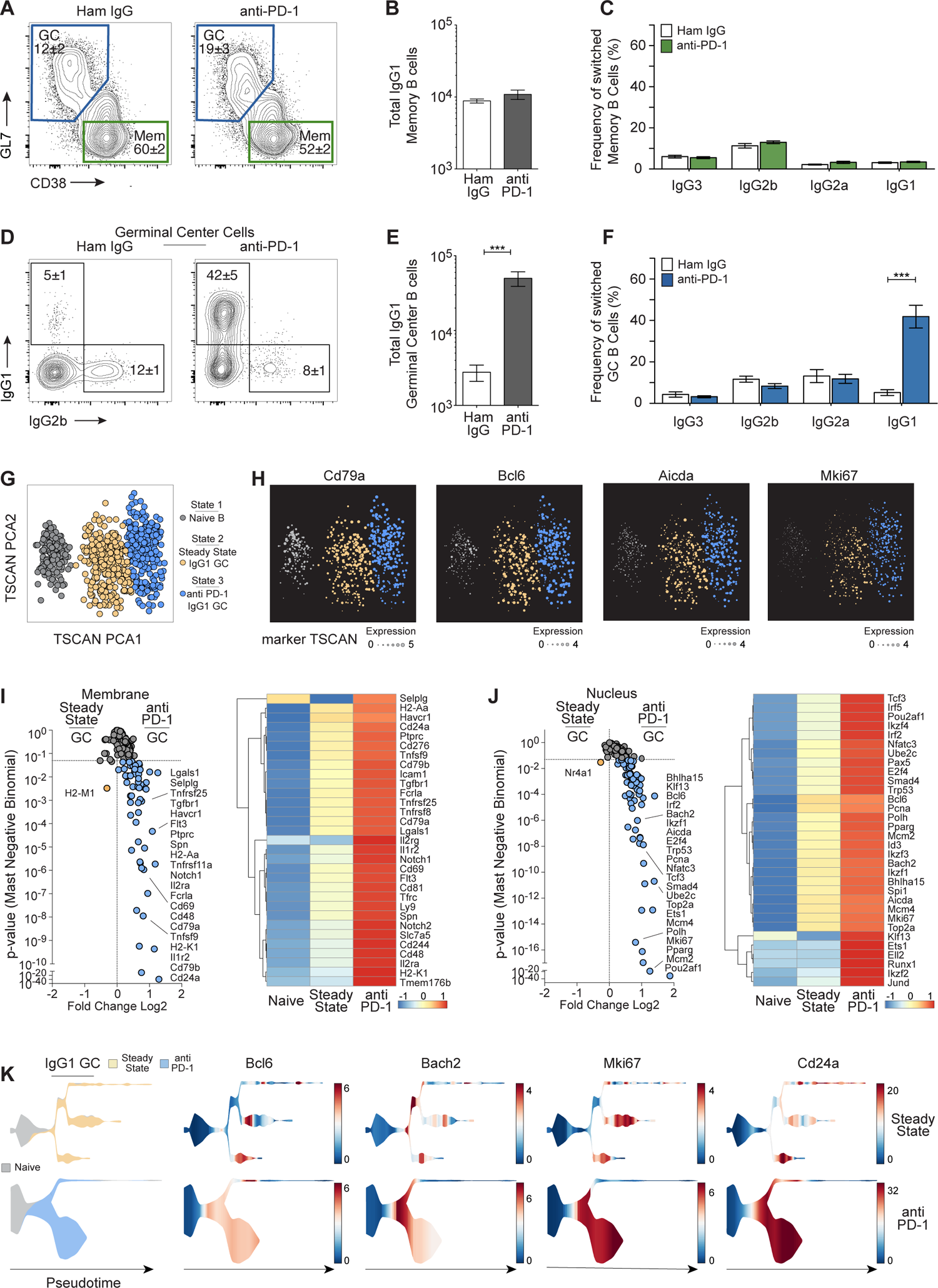
Selective expansion of IgG1 Germinal Center B cells without memory B cell involvement. **(A)** Flow cytometric analysis from mice 6 days after control IgG or anti-PD1 treatment displaying class-switched (IgD-IgM-CD138-) GC (CD38-GL7+) and memory (CD38+ GL7-) B cells, with **(B)** memory B cell quantification of IgG1, (**C)** distribution of memory IgG sub-classes and **(D)** GC distribution of IgG1 and IgG2b with **(E)** quantification for GC IgG1 (n=9) and **(F)** distribution of GC IgG sub-classes, (n=6) mean±SEM. **(G)** TSCAN pseudotime analysis of qtSEQ from single FACS sorted Naïve B cells, IgG1 GC from Steady State and anti-PD1 treated mice with **(H)** marker TSCAN gene expression. See also Figure S2. **(I)** Volcano plot of Steady State compared to anti-PD1 IgG1 GC with averaged single cell heatmap from cell membrane gene products and **(J)** nucleus. See Figure S2 for cytosol and secreted gene products and single cell heatmaps. **(K)** Stream developmental trajectories of Naïve B cells with Steady State IgG1 GC compared to anti-PD-1 IgG1 GC with sample UMI gene expression.

Similar to the IgG1^+^ PC compartment, there were substantial increases in expression for cell cycle machinery (*Mki67, Pcna; Ube2c, Trp53, E2f4*), DNA replication (*Mcm2, Mcm4, Top2a*) and DNA repair components (*Aicda, Polh*) typically associated with GC B cell expansion, somatic hypermutation and GC BCR diversification (**Figure 2J and S2H**). Clear expression increases in a multitude of TFs (*Bcl6, Bach2, Pou2af1, Id3*) highlight the induction (*Spi1, Irf2, Klf13, Bhlha15*) and orchestration of multiple complex facets (*Ikzf4, Tcf3, Ikzf3, Ets1*) of the ongoing IgG1 GC cycle after checkpoint blockade (**Figure 2J and S2H**). Using STREAM trajectory analysis (Chen et al., 2019), we display the distribution of the steady-state IgG1 GC program in contrast to the amplified and more clear segregation of GC cycle activities after treatment (*Bcl6, Bach2, Mki67, CD24a*) (**Figure 2K**). In the absence of non-GC B cell activation or memory B cell involvement, these data are consistent with the acute enhancement of a pre-existing IgG1 GC B cell cycle that skews preferentially towards IgG1 PC differentiation upon PD-1 blockade.

### Checkpoint blockade exaggerates IgG1 GC B cell programming

To further understand GC cellular dynamics, we used cell surface markers to interrogate the impact of PD-1 blockade on the zonal distribution of IgG GC B cell fate and function. The GC light zone (LZ; CXCR4^lo^CD86^hi^) B cells are selected on antigen-binding variant BCR and then with cognate TFH contact, which promotes cell cycle and BCR diversity upon re-entry into the GC dark zone (DZ; CXCR4^hi^CD86^lo^)(Victora et al., 2010). As expected, we observe that class-switched non-IgG1 GC B cells favor DZ phenotype with no change after anti-PD-1 treatment (**Figure 3A**). In contrast, IgG1 GC B cells skewed heavily into the LZ at homeostasis and then all IgG1 GC B cells expanded with a significant shift into the DZ upon PD-1 blockade (**Figure 3B**). Using single cell qtSEQ with STREAM analysis, we visualized IgG1 GC B cells with a more organized and enhanced distribution of mRNA for CD86 and CXCR4 after anti-PD1 treatment (**Figure 3C**). Using this strategy, we identified a range of highly significant divergent genes expressed in the same DZ-associated state that were heavily skewed towards cell cycle (*Pcna, Mki67*) and DNA replication and repair (*Mcm2, Mcm4, Top2a, Aicda, Polh*) released upon PD1 blockade (**Figure 3D**). Increases in expression of surface molecules (*Tnfsf9, CD24a, H2-Ab1*), signaling machinery (*Blnk, Aurkb, Dock8*) and TFs (*Pou2af1, Bcl6, Bhlha15, Bach2*) also accompanied this anti-PD1 driven amplification of DZ activity (**Figure 3D**).

**Figure 3.**
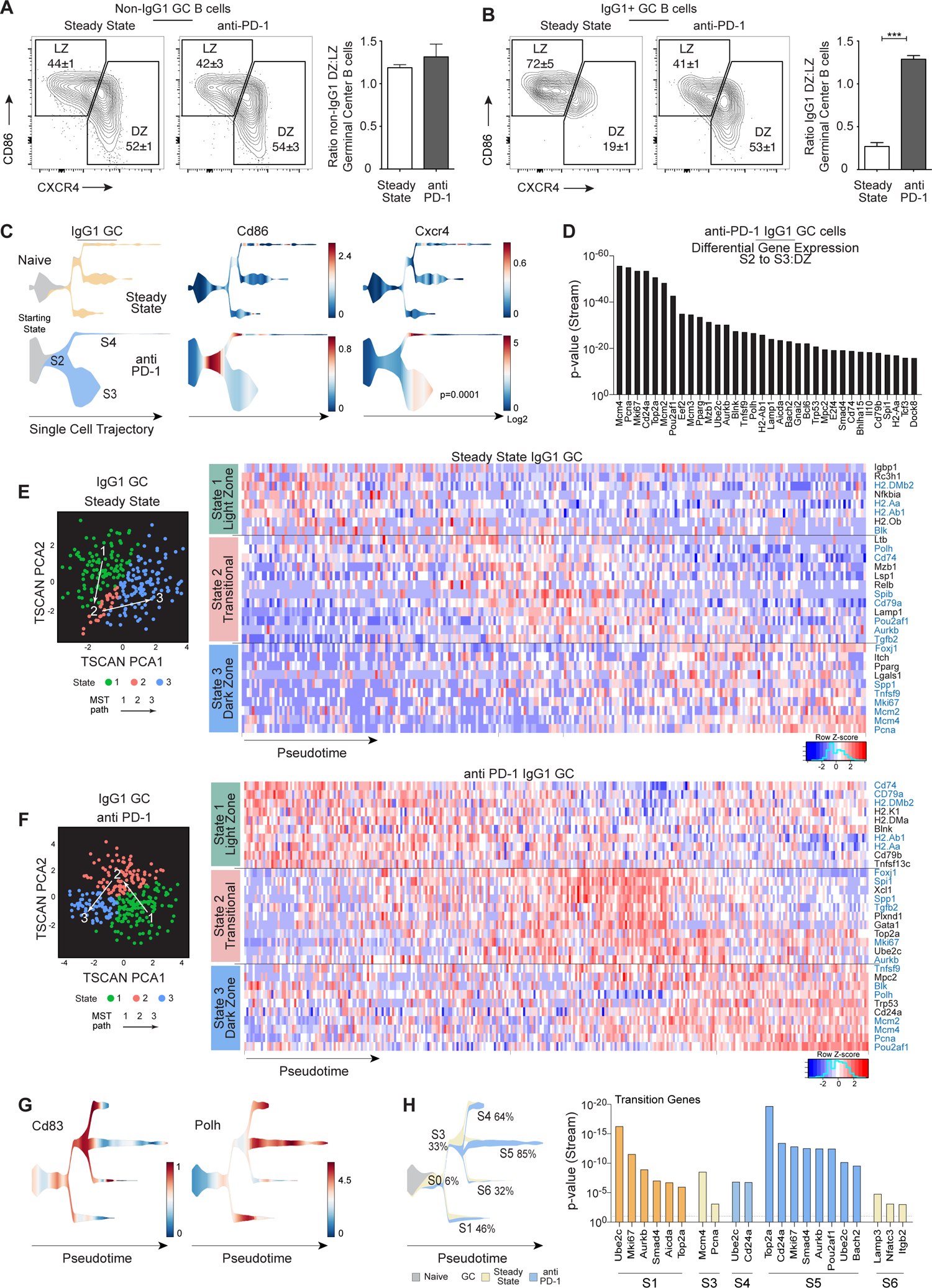
Exaggerated IgG1 GC B cell programing upon PD-1 blockade. **(A)** Flow cytometric analysis from Steady State mice or 6 days after anti-PD1 treatment displaying class-switched non-IgG1 and **(B)** IgG1 GC B cells distributed across light zone (CD86+CXCR4-) and dark zone (CD86-CXCR4+) with relative ratios (n=4) mean±SEM. (**C)** Stream developmental trajectories of single cell qtSEQ for Naïve B cells with Steady State IgG1 GC compared to anti-PD-1 IgG1 GC with distribution of *Cd86* and *Cxcr4* gene expression. Identification of starting state with diverging branches S2, S3, and S4 defined for the anti-PD-1 IgG1 GC. **(D)** Differential gene expression of S2 to S3 dark zone displaying the most significant gene products. **(E)** TSCAN pseudotime reconstruction of minimum spanning tree (MST) path and single cell heatmap displayed across each of the 3 states (State 1 LZ, State 2 Transitional, State 3 DZ) of IgG1 GC for Steady State compared to **(F)** anti-PD-1 treated mice. Gene products in blue are represented in both compartments. **(G)** Stream trajectory of combined Naïve, IgG1 GC Steady State and anti-PD1 treated mice for *Cd83* and *Polh* gene products. **(H)** Stream map with each state identified for each diverging branch (S0 to S6) and the percent contribution of anti-PD1 GC B cells, the most significant transition genes detected for each state is displayed at right.

TSCAN trajectories can be used to dissect further the organization and molecular components of anti-PD-1 driven GC cycle amplification. At steady-state, two major regions of cells displaying LZ and DZ associated activities were found with a minor separate ‘transitional’ subset (**Figure 3E**). After PD-1 blockade and IgG1 GC B cell expansion, LZ and DZ regions remained high in frequency and level of gene expression; however, a prominent ‘transitional’ zone (TZ) was discernable at equivalent frequencies to LZ and DZ subsets (**Figure 3F**). Examples of significantly divergent genes across pseudo-time are presented in heatmaps to highlight assortment in LZ-associated activities of antigen processing and presentation (*H2.Dmb2, H2-Aa, H2-Ob*), and BCR signaling (*Blk, Igbp1*) that increase (*H2.Dmb2, H2.Aa*) or change (*Cd74, Cd79b, Blnk, Tnfrsf13c, H2.K1*) upon PD-1 blockade. The range of functions released in the transitional state include notable changes in TF expression (*FoxJ1, Spi1, Gata1*), surface and secreted molecules (*Plxnd1, Spp1, Tgfb2, Xcl1*) with evidence for recent exaggerated cell cycle entry (*Aurkb, MKi67, Top2a*). These TZ enhancements are further reinforced by increases of DZ-associated gene expression for cell surface and intracellular molecules (*Tnfsf9, Cd24a, Blk*), TF expression (*Pou2af1*), and DNA replication and repair machinery (*Mcm2, Mcm4, Polh, Pcn*a, *Trp53*). Overlay STREAM reconstructions reinforce the induction of an exaggerated IgG1 GC program upon PD-1 blockade by using *Cd83* and *Polh* mRNA to discriminate LZ/DZ regions respectively (**Figure 3G**), to highlight significant changes in transition genes due to PD-1 blockade (**Figure 3H**).

Taken together, we propose that the steady-state IgG1 GC transcriptional state constitutes a tolerance program that exerts LZ constraint of the multiple pre-existing IgG1 GC B cell specificities. Acute PD-1 blockade releases this inhibitory program to exaggerate typical molecular actions of an active GC cycle that accompanies BCR maturation, ongoing cellular selection and exit into long-lived post-GC B cell compartments. Here we demonstrate the highly skewed preference towards post-GC IgG1 PC production.

### IgG1 PC transition from anti-PD-1 dysregulated GC B cell cycle

Next, we considered the molecular changes that occur between rapid IgG1 GC B cell expansion and IgG1 PC production induced by PD-1 blockade. Bray-Curtiss dissimilarity plots display high dimensional approximations of clusters for GC and PC subsets based on gene expression with further separation due to isotype and anti-PD-1 treatment (**Figure 4A**). Considering PD-1 treatment alone, TSCAN pseudo-time projections place the induced IgG1 PC program downstream from the IgG1 GC program with skewing of particular genes into IgG1 GC (*Bcl6, Aicda, Bach2*), IgG1 PC (*Xbp1, Irf4, Xbp1, Prdm1*) and across both (*Pcna, Mki67, Batf and Bhlha15*)(**Figure 4B and S3A**).

**Figure 4.**
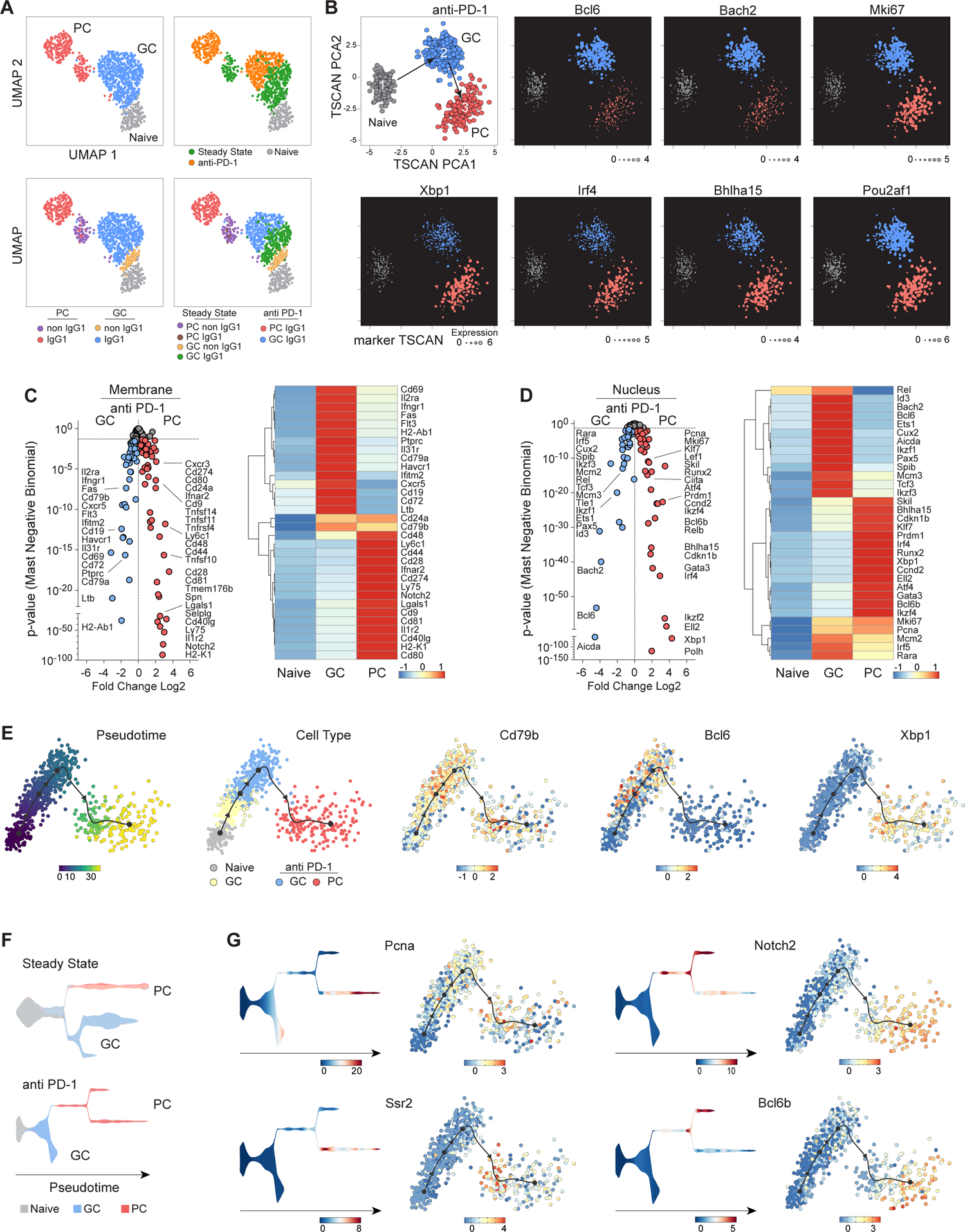
Anti-PD-1 induced transition of IgG1 GC B cell to PC program. **(A)** Single cell qtSEQ UMAP of Naïve B cell, GC and PC from Steady State and 6 days after anti-PD1 treated mice. **(B)** TSCAN pseudotime reconstruction of minimum spanning tree (MST) path for Naïve B cells and IgG1 GC and PC from anti-PD-1 treated mice with representative marker TSCAN gene expression. See also Figure S3-A. **(C)** Volcano plot comparing IgG1 GC and PC from anti-PD-1 treated mice with averaged single cell heatmap of cell membrane gene products and **(D)** nucleus. See Figure S3-B for cytosol and secreted gene products and Figure S3-C,D single cell heatmaps. **(E)** Slingshot cell lineage and pseudotime inference with sample temporal gene expression of Naïve B cell, Steady State IgG1 GC and anti-PD-1 IgG1 GC and PC. **(F)** Stream map of Naïve B cells with Steady State IgG1 GC and PC compared to Naïve B cells with anti-PD-1 IgG1 GC and PC along pseudotime. **(G)** Comparative gene expression overlaid on pseudotime distribution with Stream and Slingshot for distinguishing sample genes across PC anti-PD1 subsets. See also Figure S3-E for volcano comparison.

Closer scrutiny of these two programs reveals major differences in expressed genes with abundant state-specific and highly significant alterations that would be required to convert IgG1 GC B cells into the IgG1 PC pathway. At the cell surface, IgG1 GC B cells must lose growth factor receptors (*Il2ra, ifngr1, Flt3, Il31ra*), BCR co-receptors (*Cd79a, Cd19, Ptprc*) and T cell interactors (*Cd72, Havcr1, Cd69*), while gaining new expression of inter-cellular communicators (*Notch2, Cd48, Cd9, Lgals1*) and new means of cell contact (*Cd44, Ly75, Cd28, Cd40lg*) in this transition (**Figure 4C and S3C**). Shifts in intracellular machinery include GC decrease of BCR signaling components (*Blk, Bank1, Dock8*) and antigen processing (*H2.Dmb2, H2.Oa, Cd74*) in favor of high-fidelity protein management (Edem1, Spcs1, Fut8), efficient secretory machinery (*Mzb1,Ssr2*) with changes in metabolism (*Mpc2, Ctsz*) and secretion of soluble regulators (*Ccl21a, Il22,, Il1a, Ltbp3, Slpi*) emphasized in the IgG1 PC compartment (**Figure S3B**). Transitions in usage of major transcriptional regulators for GC (*Bcl6, Bach2, Id3, Pax5, Ets1, Ikzf1, Ikzf3, Spib, Irf5*) and PC (*Bcl6b, Bhlha15, Gata3, Ikzf4, Klf7, Atf4, Irf4, Runx2*) cohorts (**Figure 4D and S3D**) highlight the highly significant changes that are needed to drive IgG1-expressing GC B cell undergoing BCR maturation towards IgG1-secreting PC differentiation.

Using Slingshot trajectory inference (Street et al., 2018), we recreate the IgG1 GC to PC transition that accompanies PD-1 blockade and define two separable IgG1 PC subsets. We included the steady-state IgG1 GC for these pseudotime projections and separate them with overlap from the anti-PD-1 treated IgG1 GC compartments (**Figure 4E**). These data further segregate increased transcriptional activity following treatment that is restricted to the IgG1 GC (*Aicda, Bcl6, Bach2*), and extended into the PC compartment (*Pou2af1, Cd79b*) or expressed largely in the PC compartment (*Xbp1)*(**Figure 4E**). The steady-state PC compartment exists as a singular pathway while using STREAM projections, the PD-1 treated PC emerge from the GC across one node that then branches out across two forks based on transcriptional divergence (**Figure 4F**). Some genes expressed in the lower fork (*Pcna, Ssr2*) and the upper fork (*Notch2, Bcl6b*) can also be seen in slingshot projections to exemplify multiple differences in anti-PD-1 induced PC subsets (**Figure 4G and S3E**). Among many transcriptional differences, the strong indication of recent cell cycle activity with evidence of DNA replication and repair machinery upregulation is consistent with a recent GC origin, while PC population S4 has an expression profile more similar to expanded and fully differentiated IgG1 PC compartments induced by PD-1 blockade.

### PD-1 blockade provides no adjuvant action in a vaccine response

While the IgG1 selectivity has not been previously seen, the release of GC B cell control and exaggerated GC B cell expansion is predicted from previous studies of PD-1 function. Hence, we reasoned that anti-PD-1 could be used as an isotype-specific adjuvant to enhance IgG1 GC B cell maturation in the context of vaccination. We have extensive experience in the model antigen response to NP-KLH using a TLR4 agonist known to induce both IgG1 and IgG2 antibody isotypes (McHeyzer-Williams et al., 2015; Pelletier et al., 2010; Wang et al., 2012). At day 10 after priming, anti-PD-1 was introduced and shown to significantly decrease PD-1 expression on follicular T cell compartments 6 days later (**Figure S4A and S4B**). Using this strategy, we quantified the impact of acute PD-1 blockade on antigen-specific B cells from newly-formed GC (day 16 primary), persistent GC (day 48 primary) and newly-formed memory GC (day 6 recall) together with the outcomes of antigen-specific memory B cells and PC differentiation (**Figure S4C to S4I**). Surprisingly, there was no impact of PD-1 blockade on the vaccine-driven antigen-specific B cell response across all compartments and over all timepoints, regardless of antibody isotype. We thought this may be due to the overwhelming effect of the TLR4 agonist, however the pre-existing non-specific IgG1 splenic GC and PC response remained equally susceptible to PD1 blockade (at day16 and day 48 primary) but not measurable above the elevated baseline at recall (**Figure S4J and S4K**). Using high resolution flow cytometry, we can reproducibly measure even the very small antigen-specific response to NP-KLH alone without adjuvant (**Figure S4L**). Even under these sensitive circumstances with no added inflammatory drive, we saw no adjuvant impact of anti-PD1 on the resultant newly-formed antigen-specific B cell response.

### Acute checkpoint blockade releases pre-existing foreign IgG1 specificities

While PD-1 blockade did not enhance the demonstrable immune response to foreign antigen, we were able to detect a small level of pre-existing IgG1 binding to the J43 hamster antibody before treatment that assorted across all mature B cell compartments (**Figure S5A**). This specific J43 binding increased significantly over 6 days of treatment displaying the characteristic IgG1 selectivity for expansion in the GC and PC compartments (**Figure S5A**). Using ELISPOT analysis, we determined that J43 binding accounted for <30% of the total IgG1 PC expansion driven by acute blockade (**Figure S5B**). Next, we attached the foreign hapten NP to J43 under different conjugation ratios and failed to drive an anti-NP response even in the presence of the extensive anti-J43 reaction (**Figure S5C**). Taken together, we propose that PD-1 blockade does not enhance a newly-forming anti-foreign immune response, but selectively targets the release of pre-existing foreign or cross-reactive self-specificities of IgG1 isotype within the steady-state splenic GC reaction.

Broad screening for IgG1 self-specificities by immunofluorescence staining showed no detectable labeling of intracellular components from syngeneic cell lines (data not shown). However, labeling live cecal bacteria by flow cytometry captured significant increases in IgG1 binding from C57Bl/6 animals >14 weeks of age after PD-1 blockade (**Figure 5A and 5B**). To determine the specificities of IgG1 antibodies released by PD-1 blockade, we used freshly isolated cecal bacteria with detectable hamster control binding but 100-fold higher intensity binding after PD-1 blockade and sorted these bacteria for Oxford Nanopore based 16S rRNA sequencing (**Figure 5C and 5D**). Notably, there was significant enrichment in IgG1 binding to members of the order *Bacteroidales* and genus *Helicobacter*, with especially high binding to murine pathobiont species *H. typhlonius* using a steady-state CD45.1 congenic C57BL/6 microbiome containing this species (**Figure 5E**). Higher representation of binding to the genus *Lactobacillus* and species *L. johnsonii* before treatment indicates the shift away from these IgG1 specificities upon release of anti-inflammatory IgG1 antibodies with blockade. Furthermore, acute anti-PD-1 treatment significantly altered the presence of multiple gut microbial species (**Figure 5F**). *H. typhlonius* was not present in the CD45.2 steady-state C57Bl/6 microbiomes used in the latter experiments and did not emerge over the 6 days of treatment. While not significant, *Akkermansia muciniphila,* known to induce T-dependent IgG1 (Ansaldo et al., 2019), displayed the highest level of decrease over the 6 days of treatment (17-fold) in these studies. Hence, PD-1 mediated inhibition can control a subset of established anti-inflammatory anti-microbial IgG1 specificities and acute disruption significantly alters circulating antibody levels and impacts steady-state gut microbial diversity.

**Figure 5.**
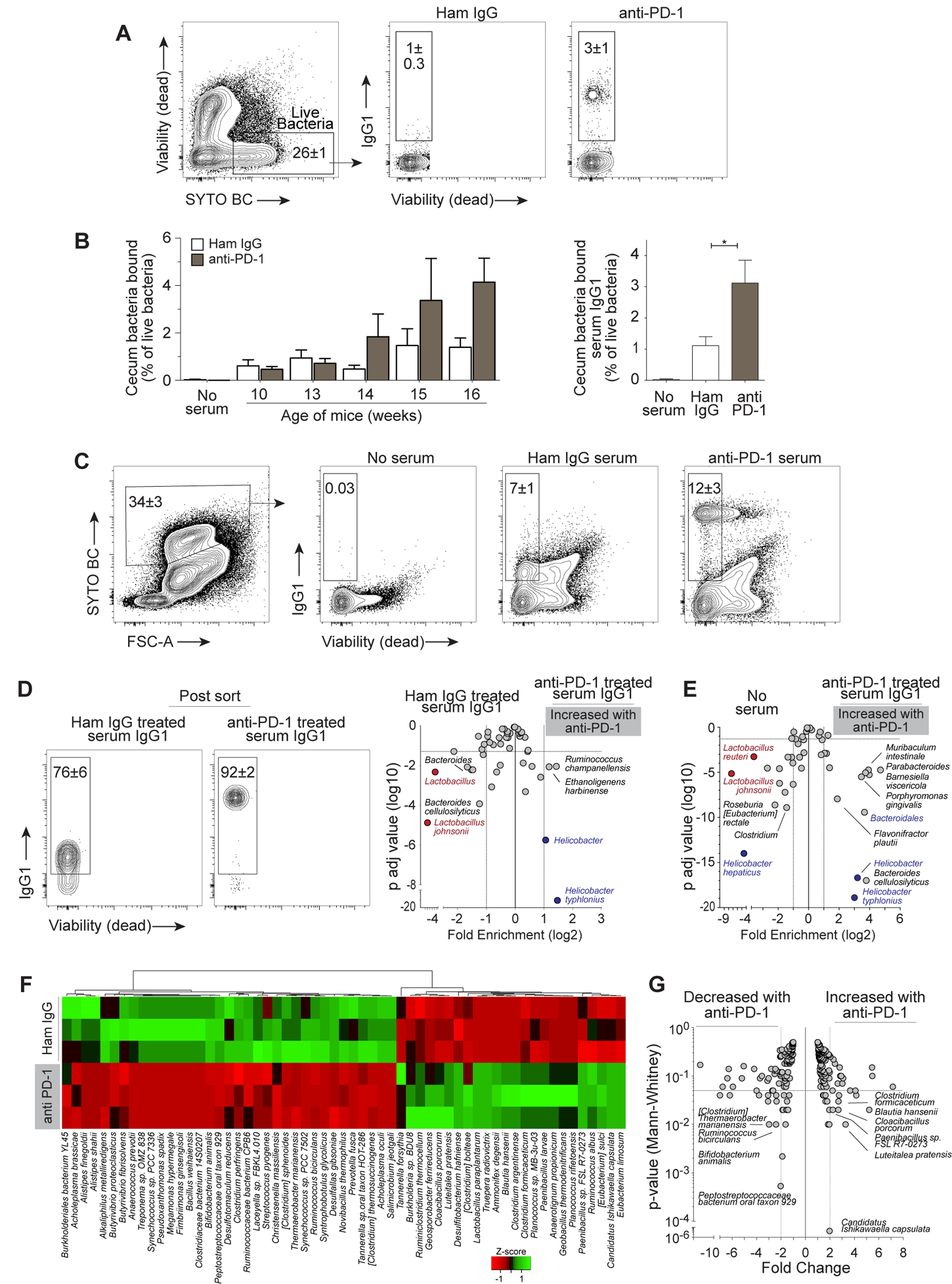
Pre-existing gut microbiota specificities released by checkpoint blockade. **(A)** Flow cytometry analysis of IgG1 binding to cecal bacteria isolated from untreated mice and incubated with serum from Hamster IgG or anti-PD-1 treated mice. **(B)** Quantification of frequency of live bacteria coated with serum IgG1 as defined in (A). Serum samples are from 5 independent experiments (n=3 for each age). Pooled data from mice 14 weeks or older (n=9). **(C)** Sorting strategy for IgG1-bound freshly isolated cecal bacteria after incubation with control serum or serum from 6 days after anti-PD-1 treatment. **(D)** Post-sort analysis and volcano plot showing comparison of serum IgG1 specificity from control-treated (n=3) to anti-PD-1 (n=4) mice or **(E)** no serum to anti-PD1 treated serum IgG1. **(F)** Heatmap displaying relative z-score for changes in species abundance and **(G)** volcano plot with fold change and significant difference (n=3).

### CD4 T cell dependent PD-1 tolerance checkpoint

The ligand PD-L1 is expressed on all memory B cell subsets and many non-IgG1 PC; however, expression is very low on all splenic GC B cells regardless of IgG subclass as well as IgG1 PC (**Figure 6A**). Hence, the selective inhibitory impact of PD-1 on IgG1 GC B cells was unlikely to involve direct signals through the PD-L1 pathway on GC B cells. Using acute depletion of CD4, we established the requirement of CD4 T cells as key regulators of the PD-1 GC B cell checkpoint (**Figure 6B**). CD4 T cell depletion did not significantly impact existing splenic IgG1 PC or memory B cell numbers; however, the enhancement of IgG1 PC by PD-1 blockade was completely abolished (**Figure 6C**). Furthermore, >95% of pre-existing IgG1 GC B cells were lost with anti-CD4 treatment, but this loss was not overcome by PD-1 blockade. These trends further reinforce the likelihood that pre-existing IgG1 GC B cells are the immediate cellular precursor of both expanded IgG1 GC B cells and post-GC IgG1 PC cell fate induced with PD-1 blockade. These data also broadly implicate CD4 T cells as the direct target of anti-PD-1 checkpoint blockade.

**Figure 6.**
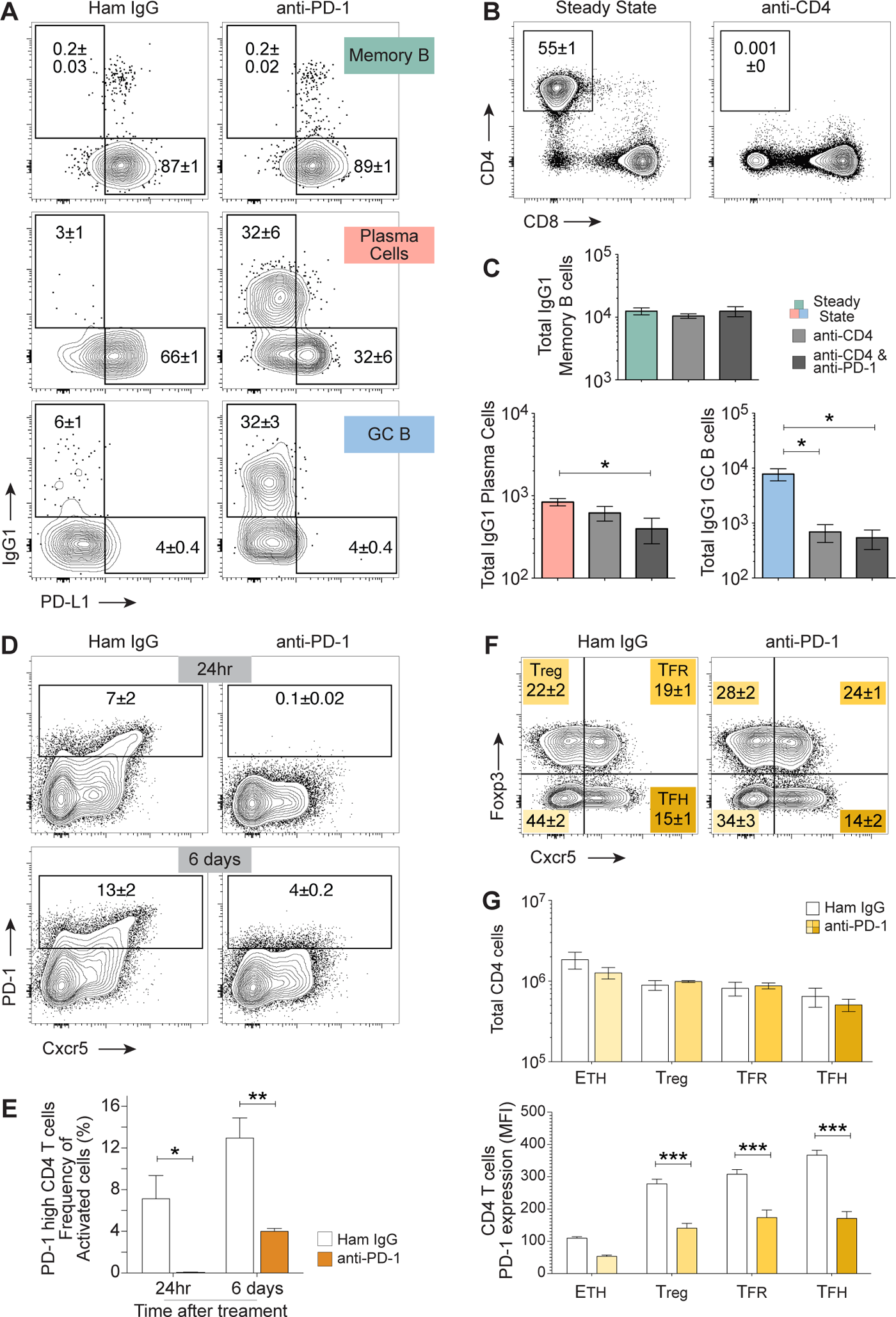
CD4 T cell dependent mechanism without local follicular T cell expansion. **(A)** Flow cytometry analysis of PD-L1 expression on splenic class-switched memory B cells, PC and GC B cells after 6 days of isotype control or treatment with anti-PD-1. **(B)** Mice treated with CD4-depleting antibody GK1.5 (anti-CD4) and **(C)** quantification of total IgG1 memory B cells, PC and GC with or without anti-PD-1 after CD4 depletion (n=3) mean±SEM. **(D)** Splenic PD-1 levels on activated CD4+ T cells (TCRb+ CD8-CD4+ CD44^hi^) following 24 hours or 6 days after control IgG or anti-PD-1 treatment with **(E)** quantification of frequency changes. **(F)** Intracellular flow cytometry of activated CD4 T cell subsets (eFluor780 viability-, GR1-, B220-TCRb+ CD8-CD4+ CD44 ^hi^) separated into major compartments after control antibody or anti-PD-1 treatment 6 days prior and **(G)** quantification of total cell numbers and PD-1 expression, (n=3) mean±SEM.

As briefly mentioned, anti-PD-1 treatment downregulates surface expression of PD-1 itself, even in the context of foreign antigen vaccination (**Figure S4B**). Here, we examined this more closely using this non-cross blocking anti-PD1 antibody with similar results at 24hr and 6 days after treatment (**Figure 6D and 6E**). In contrast to germline genetic ablation, acute PD-1 blockade had no impact on the number or broad composition of the pre-existing activated splenic CD4 T helper cell compartment at steady-state (**Figure 6F**). Even sub-fractions of conventional regulatory T cells and the two main follicular fractions of TFH and TFR cells known to express the highest levels of PD-1 show no significant alteration in numbers due to PD-1 blockade (**Figure 6G**). Hence, we propose there must be a significant acute alteration in the local transcriptional program of follicular T cells to selectively target the anti-inflammatory IgG1 pathway and release local control on IgG1 GC maturation to produce these extensive changes in the IgG1-secreting PC compartment.

## DISCUSSION

At homeostasis, the adaptive immune system remains dynamically poised between immunity and tolerance to both self and foreign antigens. Our studies reveal an acquired B cell tolerance mechanism that uses the PD-1 pathway on CD4 T cells to control B cell fate and antibody homeostasis. This PD-1 driven mechanism targets the anti-inflammatory IgG1 pathway at steady-state to constrain GC maturation and PC formation without memory B cell involvement. Using single cell qtSEQ, we define molecular components of the IgG1 GC B cell and PC tolerance mechanism and demonstrate how it is released upon acute PD-1 checkpoint blockade. We find no evidence of this activity during adjuvant-induced immunity but enhancement of pre-existing IgG1 specificities, including binding to members of the commensal microbiome, was evident upon checkpoint blockade. We propose this adaptive B cell tolerance mechanism is operative in healthy individuals to guard against over-reaction to antigens within the environment.

PD-1 blockade provides one mechanism for driving the acute progression of IgG1 GC B cells towards PC differentiation. We developed a custom qtSEQ strategy to quantify changes in the transcriptional program of individually sorted and indexed members of these rare lymphocyte sub-populations. High-fidelity cell purification is directly coupled to sensitive and targeted cDNA library formation to provide ∼10-fold increased detection per single cell of targeted mRNA species compared to global strategies (Fan et al., 2015; Gao et al., 2020; Milpied et al., 2018; Peng et al., 2021; Wu et al., 2014). PCs have been difficult to sequence due to over-abundance of Ig mRNA signals per PC impacting resolution of gene expression studies (Higgins et al., 2019; Nutt et al., 2015). Nevertheless, RNAseq strategies have defined core plasma cell programs of differentiation (Low et al., 2019; Roy et al., 2019; Shi et al., 2015), at rest and during infection in humans (King et al., 2021; Sokal et al., 2021), staged transcriptional states in the context of malignancy (Holmes et al., 2020; Ledergor et al., 2018) and implicated B cell activity and PC function in tumor microenvironments (Hollern et al., 2019; Sharonov et al., 2020). Here, we reveal acute induced alterations in intracellular machinery, changes in capacity for intercellular contact and control, and substantial shifts in key nuclear processes and TF expression that accompany IgG1 GC B cell re-programming towards PC differentiation. In the absence of IgG1 B cell activation or recruitment of IgG1 memory B cells, together with the concurrent expansion of IgG1 GC B cells, we conclude that massive local PC expansion is a directed post-GC B cell fate. Unbiased clustering and pseudo-time reconstruction provide molecular detail to progression of the IgG1 post-GC PC differentiation program that is released after acute PD-1 blockade.

Complex and progressive molecular events drive the CD4 regulated GC B cell cycle of affinity maturation (McHeyzer-Williams et al., 2012; Mesin et al., 2016; Roco et al., 2019; Victora et al., 2010; Vinuesa et al., 2016). Recent single cell molecular approaches use a range of tissue sources containing active GC to unravel the cyclic progression of GC function (Holmes et al., 2020; Kennedy et al., 2020; King et al., 2021). In the current study, the cellular distribution of splenic GC B cells suggested that the steady-state IgG1 GC B cell compartment was selectively constrained in the LZ and that PD-1 expressed on T cells was needed to restrict ongoing DZ re-entry with its associated cell cycle and BCR diversification machinery (Kennedy et al., 2020; Mesin et al., 2016; Victora et al., 2010). While the steady-state IgG1 GC B cell transcriptional program was significantly altered with treatment, it remains important to ascertain which of the many pre-existing traits were central to maintaining the adaptive tolerant state. Class-switched memory B cells and tertiary lymphoid structures (Cabrita et al., 2020; Helmink et al., 2020) have been linked to immunotherapy efficacy and patient survival (Petitprez et al., 2020) with TFH and B cell compartments also implicated (Hollern et al., 2019). In the current study, PD-1 blockade releases a vigorous and progressive IgG1 GC program with exaggerated transition towards ongoing IgG1 GC B cell DZ maturation and PC terminal differentiation. We predict that release of IgG1 GC B cell maturation can impact efficacy of checkpoint blockade in the clinic and will vary depending on the range of pre-existing GC BCR specificities in each individual.

We did not find evidence for the release of self-reactive IgG1. Here, we reveal the role of PD-1 in regulating acquired tolerance to non-self antigens (Goodnow, 2021). Unlike recent tolerance models that address the role of the GC in preventing self-reactivity (Brink and Phan, 2018; Burnett et al., 2018; Degn et al., 2017; Sabouri et al., 2014), we propose a separate mechanism driving tolerance to foreign antigen specificities encountered during normal development. This CD4 T cell-controlled tolerance mechanism constrains maturation to pre-existing GC B cell specificities to promote or permit commensalism for example. PD-1 blockade enhanced pre-existing IgG1 binding to a foreign antigen, a range of commensals with evidence for increased reactivity to a known pathobiont. We do see smaller, more variable changes to IgA GC in the Peyer’s Patch with small but significant increases in IgA PC in the bone marrow (but not spleen or PP) after acute PD-1 blockade, however the impact on the local PP IgA GC program remains uncharacterized. While IgA is more typically considered the humoral regulator of the microbiome (Donaldson et al., 2018; Fagarasan et al., 2010; Guzman-Bautista et al., 2020; Huus et al., 2021), anti-microbial IgG also impacts gut microbial composition (Chen et al., 2020; Macpherson et al., 2018; Slack et al., 2009; Yilmaz et al., 2021).

The anti-inflammatory role of murine IgG1 is similar to IgG4 in humans (Strait et al., 2015) hence restraining evolution of these GC B cell specificities offers one measure of homeostatic regulation. Checkpoint inhibitor efficacy has also been linked to microbiome composition (Gopalakrishnan et al., 2018; Matson et al., 2018; Routy et al., 2018). More recent studies connect TFH and B cell activation, together with antibody production as predictors of checkpoint inhibitor efficacy (Hollern et al., 2019). Therefore, understanding the impact of the IgG1 GC B cell checkpoint in the context of health, or its release in disease or during clinical intervention remains an important implication of our current studies.

PD-1 is highly expressed on sub-populations of CD4 TFH (Ansel et al., 1999; Fazilleau et al., 2009; Vinuesa et al., 2005) and TFR cells responsible for the proliferation and differentiation of class-switched B cells (Gonzalez-Figueroa et al., 2021; Linterman et al., 2011; Wu et al., 2020). Typically, higher levels of PD-1 are expected on the GC-localized fractions of these cells (Haynes et al., 2007; Shulman et al., 2013) and help to explain the indirect impact on GC B cells by acute blockade. Unlike genetic modification of these subsets in the germline (Good-Jacobson et al., 2010; Hams et al., 2011; Sage et al., 2013), there is no evidence of significant expansion within the steady-state TFH or TFR in the spleen in the current study. Acute ablation of CD4 T cells blocked the PD-1 enhanced IgG1 GC and PC expansion indicating the requirement of CD4s in this B cell effector phenomenon. However, we predict that the CD4 T cell program controlling IgG1 isotype-specific GC B cell and PC differentiation must be altered substantially to account for the change in adaptive B cell immune homeostasis.

The selective targeting of PD-1 blockade to the IgG1 B cell response provides one clear example of isotype-specific CD4 T cell immune regulation (Higgins et al., 2019; McHeyzer-Williams et al., 2012). We and others have proposed that TFH (Gowthaman et al., 2019; Wang et al., 2012; Zhang et al., 2020) and potentially TFR function (Gonzalez-Figueroa et al., 2021; Vinuesa et al., 2016) is organized and delivered in an isotype-specific manner. We have demonstrated the IgG2a-selective memory B cell requirement for T-bet expression, not only for class-switch, but also for survival into memory and re-expansion at antigen recall (Wang et al., 2012) and attribute this requirement to differential memory TFH organization. On the T cell perspective, Eisenbarth and colleagues (Gowthaman et al., 2019) recently discovered a TFH subtype with a selective role in controlling the IgE allergic. Here, we isolate the IgG1 pathway as susceptible to PD-1 blockade with further preference for GC function and PC formation without memory B cell involvement. The murine IgG1 isotype is known to be anti-inflammatory (Strait et al., 2015), induced by the same CD4 T cell signals that drive IgE class switch (Snapper and Paul, 1987; Yang et al., 2020) and importantly, IgG1 GC B cells are considered the ‘reservoir’ B cell state immediately upstream of IgE-secreting PCs (He et al., 2017; Turqueti-Neves et al., 2015). Thus, we propose that this PD-1 adaptive B cell tolerance checkpoint operates as an antigen-specific rheostat for IgG1 GC B cell maturation that prevents hypersensitivity to the commensal microbiome and other environmental antigens.

Our observations identify PD-1 as a key tolerance checkpoint in the adaptive IgG1 isotype-specific GC B cell to PC axis of differentiation. We found no consistent evidence of dysregulated IgE GC or PC production in our studies or extending into Balb/c animals that favor IgE responses. Hence, the broader connection to allergy must be explored with the notion that a second checkpoint exists for the separate and subsequent induction of IgE producing antibodies in allergy. We provide evidence for an IgG1-specific tolerance mechanism and how it is released upon checkpoint blockade across both states of IgG1 GC B cell and IgG1 PC based immunity. We propose molecular links between these functionally distinct but temporally related B cell immune fate and functions. The molecular targets revealed could prove effective for clinical modification at times of checkpoint blockade or otherwise. It will be important to extend these new principles and further studies into humans at health, disease and following checkpoint blockade.

## ACKNOWLEDGEMENTS

Graphical Abstract constructed using BioRender.com. This work was supported by the US National Institutes of Health (AI047231, AI040215 and AI071182) and Bill & Melinda Gates Foundation (BMGF 0PP1154835) to M.G.M.-W.

## AUTHOR CONTRIBUTIONS

Conceptualization: C.R.D., L.M.W., M.M.W.; Design and Methodology: C.R.D., A.G.S., B.W.H., L.M.W., M.M.W.; Acquisition, Analysis and Interpretation of Data: C.R.D., A.G.S., B.W.H., L.M.W., M.M.W.; Writing and Review of manuscript: C.R.D., A.G.S., B.W.H., L.M.W., M.M.W.; Supervision: L.M.W., M.M.W.; Funding Acquisition: M.M.W

## EXPERIMENTAL MODEL DETAILS

### Mice

Male 8-18 week old C57BL/6 and congenic C57BL/6-Ly5.1 (B6.SJL-Ptprc^a^ Pepc^b^/BoyJ) mice were bred and housed under specific-pathogen-free (SPF) conditions at The Scripps Research Institute. Mice were euthanized with isoflurane prior to sample collection and analysis. All experiments were performed with protocols approved by Institutional Animal Care and Use Committee at The Scripps Research Institute.

## METHOD DETAILS

### In vivo Antibody treatments & Immunization

For in vivo blockade of PD-1, 300µg anti-mouse PD-1 antibody (clone J43, 29F.1A12 or RMP1-14, BioXcell) was administered intraperitoneal (IP). Control mice were given 300µg polyclonal Armenian hamster IgG, or rat IgG2a isotype control, anti-trinitrophenol (clone 2A3, BioXcell). For CD4 depletion, 300µg anti-mouse CD4, GK1.5 monoclonal antibody (BioXcell) was administered. Mice were analyzed 6 days (or as indicated) post-blockade treatment.

The hapten 4-hydroxy-3-nitrophenylacetyl (NP, Biosearch) was conjugated to keyhole limpet hemocyanin (KLH, Pierce; NP-KLH, in house preparation, (Baldridge and Crane, 1999; McHeyzer-Williams et al., 1991), or anti-PD-1 antibody clone J43 (BioXcell; NP-J43). Immunizations were performed by subcutaneous (sc) or IP injection of 300 or 400µg immunogen with & without Monophosphoryl lipid A (MPL)-based adjuvant (2% squalene, 0.5 mg MPL, in house preparation,(Baldridge and Crane, 1999), as indicated.

### Flow cytometry

Single cell suspensions from spleen were prepared (Wang et al., 2012). After lysing with ACK red cell lysis buffer (Lonza) and passed through a 70 µm cell strainer (Fisher Scientific), cells were washed and resuspended at 4 x 10^8^ cells/mL in FACS buffer: 2% FCS, 10 mM HEPES in phenol red-free DMEM with L-glutamine (Gibco). Draining lymph nodes for base of tail immunization (inguinal and periaortic lymph nodes, LN) were combined and processed together. All processing and staining were performed on ice. Prior to staining, Fc receptors were blocked with 10 µg/mL of purified antibody clone 2.4G2 (BioXcell) for 10 min.

For surface staining, a master mix containing a panel of fluorophore- or biotin-labeled antibodies was mixed at pre-titrated concentrations in Brilliant Violet Staining Buffer (BD Biosciences) at 2X, and added to suspended cells at a 1:1 ratio for a final staining concentration of 2 x 10^8^ cells/mL for 30-45 min. For secondary staining of biotin-labeled antibodies, cells were washed and resuspended in FACS buffer. Fluorophore-conjugated streptavidin was added for 15-30 min. Cells were washed and resuspended in FACS buffer before analysis.

For panels including surface staining of IgG, cells were first stained with single isotype-specific fluorophore- or biotin-conjugated antibodies for 30 min. After washing and resuspending in FACS buffer, cells were blocked with 1% normal rat serum and 1% normal mouse serum (Jackson ImmunoResearch) for 10 min. Next, staining using the master mix of additional antibodies and streptavidin reagents was performed as above. If the isotype-specific antibody was biotin-conjugated, the fluorophore-conjugated streptavidin was included with the master mix stain.

For intracellular FACS, surface stains were completed as described above. Cells were incubated with eBioscience Fixable Viability Dye eFluor 780 (Invitrogen) for 10 min, washed with FACS buffer, then fixed and permeabilized using the eBioscience Foxp3 / Transcription Factor Staining Buffer Set (Invitrogen). Cells were incubated in 1X Fixation/Permeabilization buffer for 25 min and washed with 1X Permeabilization buffer. Prior to intracellular staining, nonspecific binding was blocked by incubation with 1% normal mouse serum and 1% normal rat serum for 10 min. Cells were then stained with fluorophore-conjugated antibodies specific for intracellular antigens. Finally, cells were washed once with 1X Permeabilization buffer, and once with FACS buffer prior to resuspension in FACS buffer for analysis on the flow cytometer.

All flow cytometry sample data was collected using a BD FACSAria III on a workstation running FACSDiva software version 8.0.1 (BD Biosciences). Cytometry data analysis was performed using FlowJo version 10 (TreeStar). FACS data is presented with the following gates: plasma cells (Gr1-CD3e-IgD-IgM-CD138+), germinal center B cells (Gr1-CD3e-IgD-IgM-CD138-CD19+B220+CD38-GL7+), memory B cells (Gr1-CD3e-IgD-IgM-CD138-CD19+B220+GL7-CD38+), NP-specific B cells (Gr1-CD3e-CD19+ and/or CD138+IgD-IgM-NP+Lambda1+), activated CD4 T cells (eFluor780 Viability-Gr1-B220-TCRβ+CD8-CD4+CD44^hi^).

### Elispot

As previously described (Pelletier et al., 2010), multiScreen-HA sterile 0.45 µm surfactant-free mixed cellulose ester membrane filter plate (Millipore) wells were coated with 50 µl of 25 µg/mL unlabeled goat anti-mouse IgG (SouthernBiotech), anti-mouse IgG1 (RMG1-1, BioLegend), polyclonal Hamster IgG (BioXcell), or J43 (BioXcell), for 4 hours at room temperature or overnight at 4°C. The plate was washed 3 times with sterile PBS prior to plating samples. Cells were resuspended in RPMI medium with L-glutamine (Gibco) containing 5% fetal calf serum (FCS), 10 mM HEPES, 100 units/mL penicillin, and 100 µg/mL streptomycin, and plated in two-fold dilutions. Negative control wells were included, for which medium only was plated. After incubation overnight at 37°C and 5% CO_2_, the plate was washed 3 times with 0.1% Tween 20 in PBS and 3 times with PBS, then incubated with horse radish peroxidase (HRP) conjugated goat anti-mouse Fc-specific antibodies in FACS buffer with 5% FCS and 10% dry instant nonfat milk. The plate was incubated for 4 hours at room temperature, washed as in the previous step, and developed with chromogen substrate: 0.03% 3-amino-9-ethylcarbazole (Sigma-Aldrich) and 0.015% H_2_O_2_ in 1M sodium acetate, pH 4.8-5.0. Spots were enumerated under a dissection microscope, and plasma cell numbers were determined by extrapolation from total cell counts. All positive wells containing discernible spots were averaged together to obtain per-sample final counts.

### FACS analysis of microbiome-specific serum Ig

Bacterial flow cytometry was performed as previously described (Bunker et al., 2017). Cecum contents were squeezed into a sterile 2 mL tube on ice with 1 mL FACS Buffer and homogenized by vortexing horizontally for 5 min at 3,000 rpm. After centrifugation for 400 x g for 5 min to eliminate large debris, the supernatant was centrifuged once again at 8000 x g for 5 min to pellet cecum bacteria. The supernatant was then aspirated, and the bacterial pellet washed twice with FACS buffer. The washed bacteria were resuspended in 1 mL FACS buffer and quantified by taking the optical density at 600 nm using the DU 530 Life Science UV/Vis Spectrophotometer (Beckman). Bacterial concentration was determined assuming that:

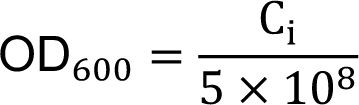

where C_#_ represents the concentration in bacteria/mL. Bacteria were resuspended in FACS buffer and all subsequent staining steps were performed at 2 x 10^8^ cells/mL. All processing was performed on ice.

Bacteria were first stained with eBioscience Fixable Viability Dye eFluor 780 (Invitrogen) and STYO BC Green Fluorescent Nucleic Acid Stain (Invitrogen) at 1:750 dilution each for 30 min. Bacteria were washed with FACS buffer and centrifuged at 8,000 x g for 5 min. To assess serum Ig binding, bacteria were incubated with two-fold serial dilutions of serum, beginning with a 1:12.5 dilution to a 1:X dilution, for 45 min. Bacteria were washed and stained with phycoerythrin (PE)-conjugated anti-IgG1 antibody for 20 min. After a final wash step, bacteria were resuspended in FACS buffer and analyzed using a FACSAria III flow cytometer.

To sort bacteria for downstream 16S rRNA sequencing, bacteria were sorted into 5 mL tubes containing 500 µl DMEM (phenol red-free, no antibiotics). Sorted samples were centrifuged at 13,000 x g for 10 min. The supernatant was aspirated, and the pellet was frozen at −80°C until genomic DNA library preparation (at least overnight).

### 16S rRNA sequencing analysis

Genomic DNA was purified from sorted bacteria, or raw cecum or feces. Raw cecum or feces was frozen for at least overnight at −80°C before DNA purification. Bacterial DNA was purified using the DNeasy PowerSoil kit (Qiagen) as per manufacturer’s instructions, under sterile conditions. DNA was eluted off the spin column with sterile nuclease-free water (Teknova) and quantified using the Qubit dsDNA High Sensitivity Assay Kit (Invitrogen). A microbial community standard (Zymo Research) was included in genomic DNA purification steps and subsequent library preparation and analysis.

16S rRNA library preparation was performed using the 16S Barcoding Kit from Oxford Nanopore Technologies (#SQK-RAB204), as per manufacturer’s instructions. Twelve samples including the microbial community standard and a reagent-only control were processed separately in parallel. Briefly, 10 ng of genomic DNA was amplified using LongAmp Taq 2X Master Mix (New England BioLabs) and the provided barcoded 16S primers. PCR was performed using an initial denaturation for 1 min at 95°C, followed by 25-30 cycles of: 20 sec at 95°C, 30 sec at 55°C, and 2 min at 65°C, and a final extension of 5 min at 65°C. Cleanup of PCR product was performed using AMPure XP beads (Beckman Coulter), and the concentration of amplified DNA for each sample was measured using the Qubit dsDNA HS Assay Kit as above. Assuming a PCR product length of 1500Kb, all samples were pooled in roughly equal molar proportion. A MinION Flow Cell version R9.4.1 was prepared for 1D sequencing as per manufacturer’s instructions, and a total amount 50-100 fmol of DNA was sequenced using the MinION Mk1B and MinKNOW (Oxford Nanopore Technologies). Fast5 files generated over the course of the 48-hour sequencing run were transferred to a computing cluster for downstream basecalling and analysis.

Pre-processing of reads collected using MinKNOW version 3.1.19 was performed using Albacore version 2.3.3 (Oxford Nanopore Technologies) and custom scripts to basecall, de-barcode, and convert to FASTQ separate directories of 40,000 reads in parallel. These FASTQ files were concatenated by barcode and trimmed using Porechop version 0.2.3 (https://github.com/rrwick/Porechop) with the default options selected.

Pre-processing of reads collected using MinKNOW version 3.1.19 was performed using the guppy_basecaller module of Guppy version 2.3.5 (Oxford Nanopore Technologies) using the following options: qscore_filtering, min_qscore 7, calib_detect, records_per_fast 0. De-barcoding was performed separately using the guppy_barcoder module of Guppy, specifying the SQK-RAB204 kit and -q value of 0.

Alignment of basecalled, trimmed, and de-barcoded reads was performed using Kraken version 2.0.7-beta (https://ccb.jhu.edu/software/kraken2). The Kraken 2 reference library used for sequence assignment was built from Archaea and Bacteria sequences downloaded from the NCBI database on August 12, 2018, using the default settings. Sequence assignment reports were converted to BIOM format (McDonald et al., 2012) using kraken-biom version 1.0.1 (https://github.com/smdabdoub/kraken-biom) and then converted to a DESeq object using the phyloseq R package version 1.29.0 (McMurdie and Holmes, 2013). DESeq2 was used to calculate differential representation of assigned taxa between groups, using the default multiple-inference correction (Benjamini-Hochberg). Prior to analysis, geometric means of counts were used to estimate size factors for normalization. Expression-based heatmap visualization was preformed using Heatmapper (Babicki et al., 2016) (http://www.heatmapper.ca) with average linkage clustering and Pearson distance measurement.

### Integrated Single Cell RNA-seq (qtSEQ)

Single cell transcriptional analysis was carried out using a custom RNA sequencing protocol called quantitative targeted single cell sequencing (qtSEQ). This technique sequences polyA mRNA using a nested set of 3’ targeted primers to amplify specific gene sets.

Single naïve B cells (Gr1-FceR1a-IgD+IgM^low^CD138-GL7-CD79b+CD19+CD38+), plasma cells (Gr1-FceR1a-IgD-IgM-CD138+IgG1+), germinal center cells (Gr1-FceR1a-IgD-IgM-CD138-CD79b+CD19+CD38-GL7+IgG1+) from 8 individual mice were sorted into 384 well plates at 4C. Doublets were excluded using SSC-A vs SSC-H. Each well contained 1µL of reverse transcription (RT) master mix consisting of 0.03μL SuperScript II taq, 0.2μL SuperScript II 5x buffer, 0.012μL of 25μM dNTPs, 0.03μL DTT (0.1M stock), 0.03μL RNaseOUT (Invitrogen), 0.19μL DNase/RNase Free Water and 0.5μL of extended oligoDT primers (1μM stock, custom made, IDT). To allow assessment of contamination from sample processing, 8 negative control well with no cell sorted were interspersed across each plate. Immediately after sorting, reverse transcription (RT) was performed at 42°C for 50 mins then at 80°C for 10 mins.

Following RT, all 384 wells were pooled within each plate and excess oligoDT primer removed using ExoSAP-IT Express (Affymetrix) at 37C for 20mins. cDNA was purified and volume reduced using a 0.8x AMpure XP SPRIselect magnetic bead (BeckmanCoulter) solution to 1x library ratio.

For each plate two nested PCR reactions were used to amplify specific gene targets then a final PCR reaction perfomed to attach Illumina sequencing adapters. Library preparation was carried out on each plate separately to preserve cell labeling.

PCR 1 was performed in a 60μL reaction: 35μL cDNA library in H2O, 1.5μL RNaseH (5000U/mL, NEB), 1.2μL Phusion DNA polymerase (2000U/mL, NEB), 12μL 5X Phusion HF Buffer (NEB), 0.85μL of 25μM dNTPS (Invitrogen), 1.2μL of 100μM RA5-overhang primer (custom made, 5’AAGCAGTGGTGAGTTCTACAGTCCGACGATC 3’) and 25nM final concentration for each specific gene targeting external primer using the following thermocycling conditions: 37°C for 20 minutes, 98°C for 3 minutes, followed by 10 cycles of 95°C for 30 seconds, 60°C for 3 minutes, 72 °C for 1 minute and final elongation at 72°C for 5 minutes. Removal of excess primers and volume reduction was performed again using 0.9x AMpure XP SPRIselect beads to 1x library ratio.

PCR2 was performed in a 20μL reaction: 10μL PCR1 reaction elute, 0.4μL Phusion DNA polymerase (2000U/mL, NEB), 4μL 5X Phusion Buffer (NEB), 0.3μL of 25μM dNTPs (Invitrogen), 2μL of 20μM RA5 (custom made, 5’ GAGTTCTACAGTCCGACGATC 3’), and 25nM final concentration for each specific gene targeting internal primer using the following thermocycling conditions: 95°C for 3 minutes followed by 10 cycles of 95°C for 30 seconds, 60°C for 3 minutes, 72 °C for 1 minute and final elongation at 72°C for 5 minutes. Removal of excess primers and volume reduction was performed using AMpure XP SPRIselect beads at a 0.9x bead to 1x library ratio.

PCR3 was performed in a 20μL reaction: 10μL PCR2 reaction elute, 0.4μL Phusion DNA polymerase (2000U/mL, NEB), 4μL 5X Phusion Buffer (NEB), 0.3μL of 25μM dNTPs (Invitrogen), 2μL of 10μM RP1 primer (Illumina), 2μL of 10μM RPI library specific primer (Illumina) and 1.3μL water using the following thermocycling conditions: 95°C for 3 minutes followed by 8 cycles of 95°C for 15 seconds, 60°C for 30 seconds, 72 °C for 30 seconds and final elongation at 72°C for 5 minutes. Removal of excess primers and volume reduction was performed using 0.7x AMpure XP SPRIselect beads to 1x library ratio and then eluted in 20μL H2O for quantification and library sequencing preparation.

Final cDNA amplified library concentration and quantity was determined using a Qubit 4 Fluorometer (Invitrogen Technologies) and 2100 BioAnalyzer (Agilent). Separately labeled cDNA libraries were mutliplexed according to manufacturer’s protocol (Illumina, San Diego CA). Libraries were sequenced using an Illumina NextSeq 500 (Read 1: 19 cycles; Index Read 1: 6 cycles; Read 2: 67 cycles).

## QUANTIFICATION AND STATISTICAL ANALYSIS

### Data Processing and Normalization

Bcl2fastq (Illumina) was used to demultiplex sequencing libraries using plate barcodes and assort reads by well barcodes prior to sequence alignment using Bowtie2 (v2.2.9) to a custom genome consisting of the amplicon regions for the specifically targeted genes used in the experiment based on the murine genome GRCm38.p6, version R97 (Ensembl). HTseq was used for read and UMI reduced tabulation. Custom scripts for these steps available upon request.

Quality control excluded cells that contained fewer than 24 UMI counts and had fewer than 14 unique genes detected. A secondary quality control filter included naïve B cells that contained 2 of 3 genes (*Cd79a, Cd79b, Ptprc*) n=158, germinal center cells containing 2 of 5 genes (*Cd79a, Cd79b, Aicda, Bcl6, Fas*) n=623, and plasma cells containing 2 of 5 genes (*Cd79a, Cd79b, Prdm1, Irf4, Xbp1*) n=266. A total 1047 B cells from 2 experiments were analyzed.

Data from the two separate experiments was pooled into a single count matrix for processing using the R package Seurat (v3.2.3) (Stuart et al., 2019); scaling and normalization was carried out separately by cell type using the ScTransform package (v0.3.2.9002, (Hafemeister and Satija, 2019) with the batch_var option set to the metadata variable “Experiment”. This normalized UMI count matrix was then used for downstream analysis.

### scRNA Sequencing Computational Analysis

Using the gene expression UMI normalized count matrices, Tools for Single Cell Analysis (TSCAN) (v1.0, (Ji and Ji, 2016) constructed cluster centroids, minimum spanning tree (MST) pseudo-temporal path, identified underlying marker genes and heatmap expression of top significant genes (FDR p<0.05) that increased along each pseudotime MST path. For the heatmaps generated, each column represents a cell that was mapped and ordered by its pseudotime value with each row denoting significant genes along the MST path.

For differential gene expression analyses, MAST (Model-based Analysis of Single Cell Transcriptomics) package (v1.16.0 https://github.com/RGLab/MAST/ (Finak et al., 2015)) using a hurdle model was used to generated fold change and p-values for all genes. Volcano plots were then created using Prism v9.0 (GraphPad Software). The R package pheatmap (v1.0.12) was used to plot heatmaps of averaged UMI counts with euclidean distance hierarchical clustering. Single cell heatmaps were generated using the “Doheatmap” function from the Seurat package.

Normalized UMI count matrices were used for psuedotime trajectory analysis with STREAM (Single-cell Trajectories Reconstruction, Exploration And Mapping of omics data)(Chen et al., 2019)(https://stream.pinellolab.partners.org/) and Slingshot (Street et al., 2018). Slingshot trajectory analysis was performed using the “dyno” analysis package (https://github.com/dynverse/dyno).

Uniform Manifold Approximation and Projection (UMAP) was performed using Biovinci (v3.0.9 Bioturing) on all genes. Statistical tests were calculated and graphed using Prism v9.0 (GraphPad Software). A P value *<0.05, **<0.01, ***<0.001 was considered statistically significant.

---

**Figure S1.**
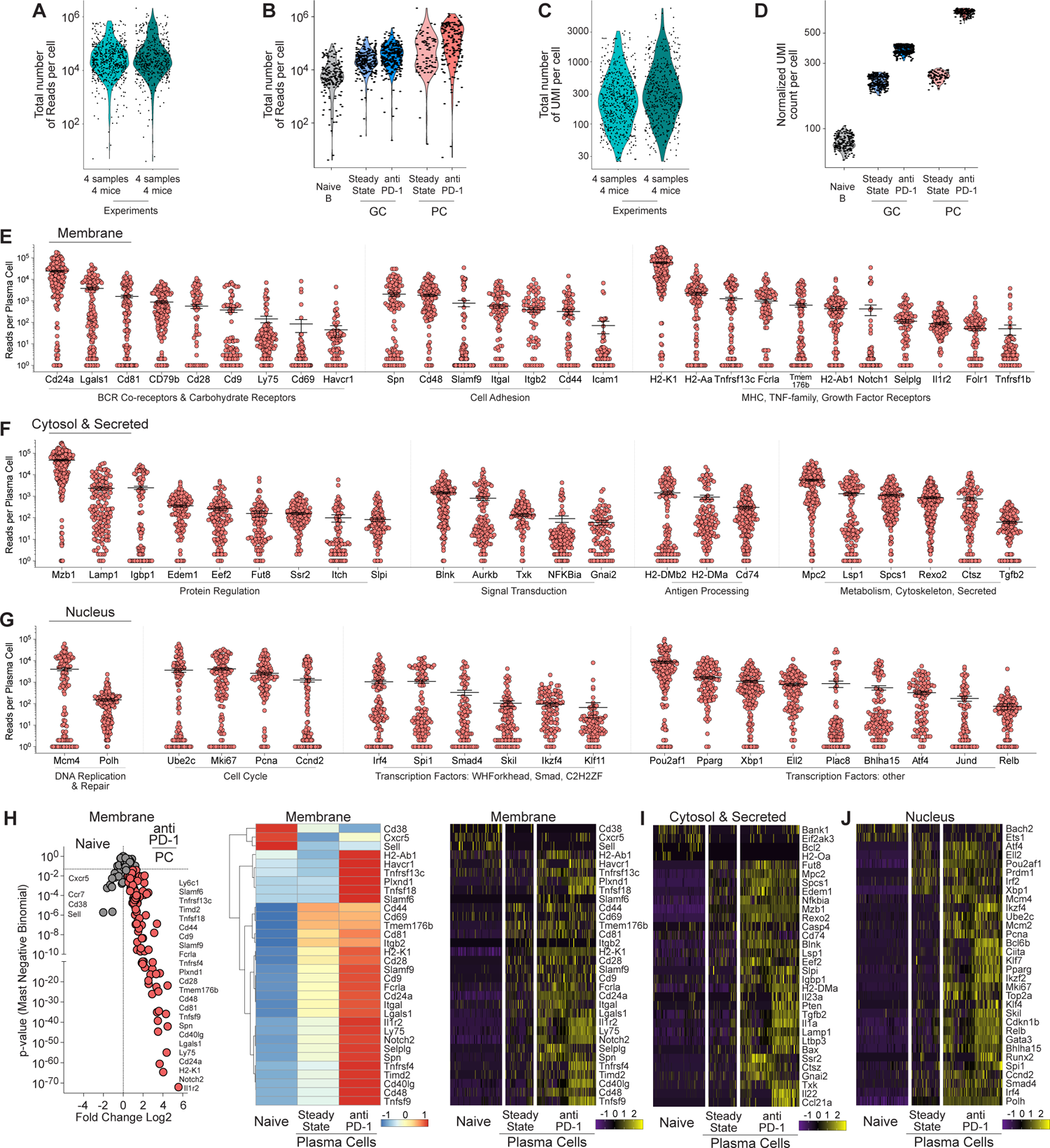
Integrated single cell RNA-seq analysis (qtSEQ) of Plasma Cell differentiation after PD-1 blockade, Related to. Figure 1 **(A)** Single-cell RNA-sequencing with qtSEQ violin plots showing total reads per cell from sorted cells that passed quality control divided across experiments and **(B)** cell type. **(C)** Total UMI counts per cell across experiments and **(D)** cell type after normalization using SCTransform. **(E)** Reads per cell from sorted anti-PD-1 IgG1 PCs for selected assays targeting gene products from cell membrane, **(F)** cytosol and secreted, and **(G)** nucleus. **(H)** Volcano plot of Naïve B cells compared to anti-PD1 IgG1 PC with averaged single cell heatmap (including Steady State PC) and single cell heatmap from UMI counts for cell membrane gene products, **(I)** cytosol and secreted and **(J)** nucleus.

**Figure S2.**
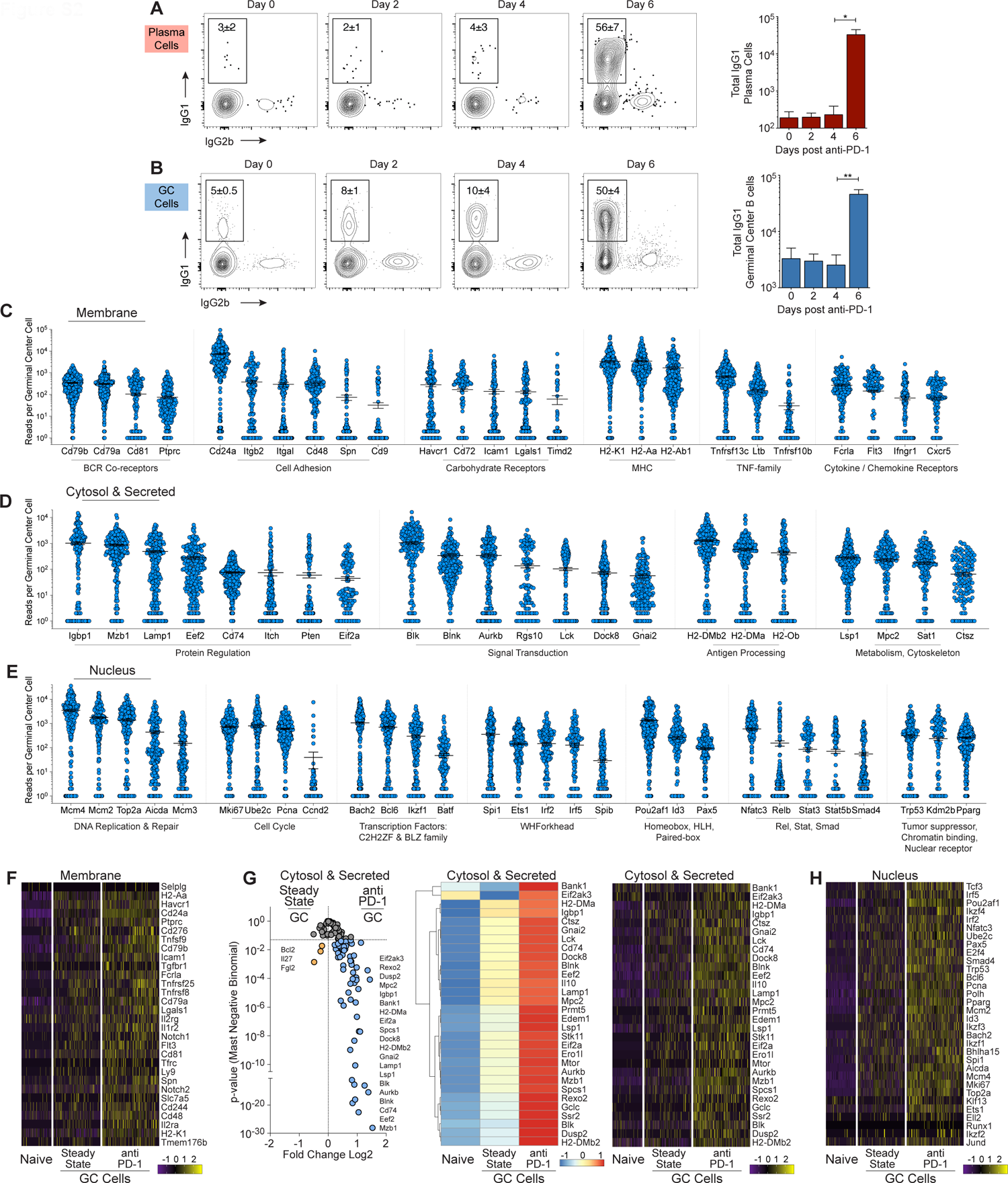
Kinetics of PD-1 blockade and single cell qtSEQ of the Germinal Center, Related to. Figure 2 **(A)** Timecourse analysis of mice either untreated (day 0) or treated with anti-PD-1 for splenic PC and **(B)** GC B cells at various days with quantification of total IgG1 cells (n=3) mean±SEM. **(C)** Reads per cell from sorted anti-PD-1 IgG1 GC for selected assays targeting gene products from cell membrane, **(D)** cytosol and secreted, and **(E)** nucleus. **(F)** Single cell heatmap from UMI counts for cell membrane gene products and **(G)** Volcano plot of Steady State compared to anti-PD1 IgG1 GC with averaged single cell heatmap and single cell heatmap from cytosol and secreted gene products. **(H)** Single cell heatmap from gene products from nucleus.

**Figure S3.**
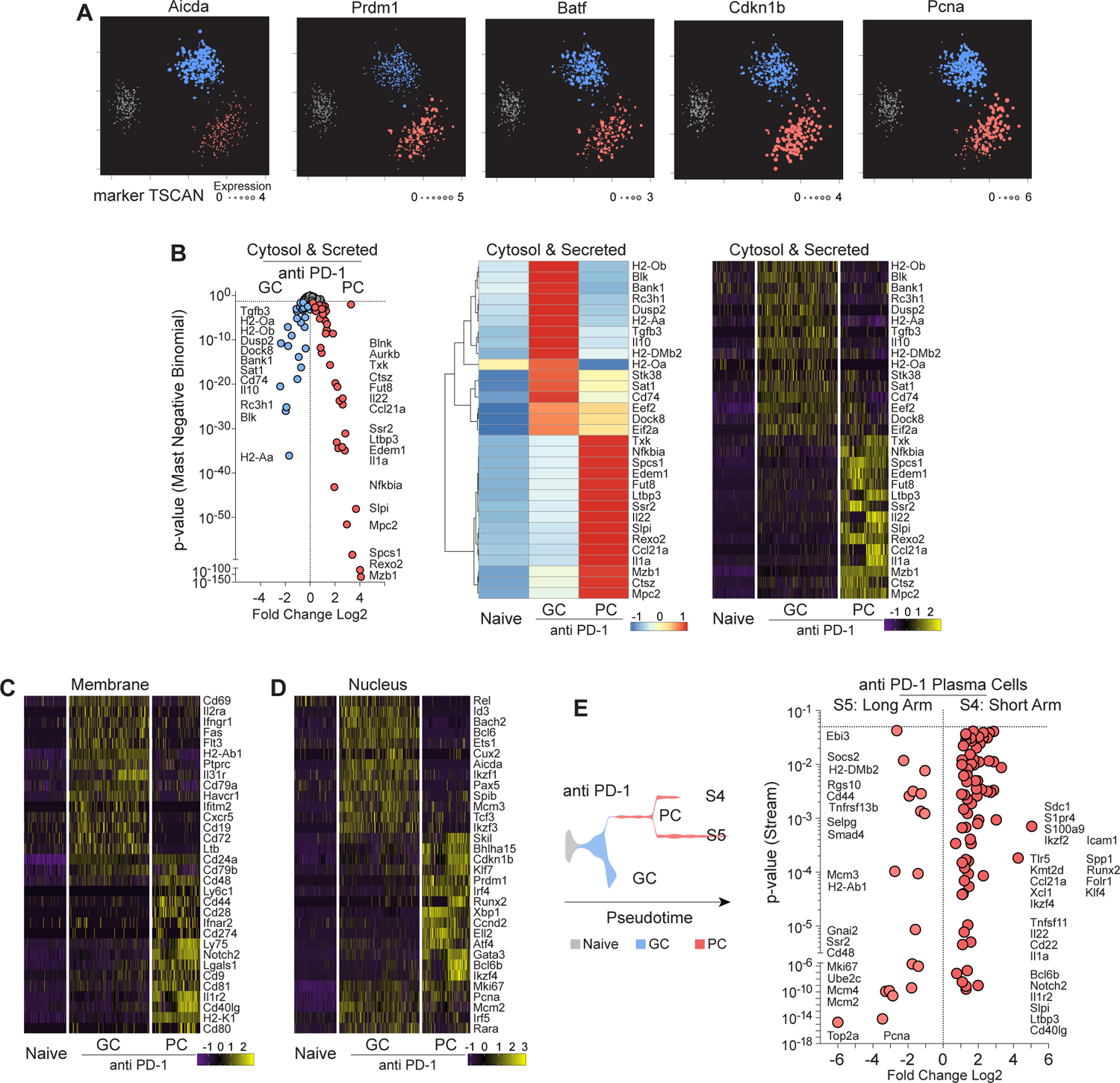
Single cell qtSEQ comparison of GC to PC program after PD-1 blockade, Related to. Figure 4 **(A)** TSCAN pseudotime reconstruction for Naïve B cells and IgG1 GC and PC from anti-PD-1 treated mice with representative marker TSCAN gene expression. **(B)** Volcano plot of IgG1 GC and PC from anti-PD-1 treated mice with averaged single cell heatmap from cytosol and secreted gene products and the single cell heatmap. **(C)** Single cell heatmap of gene products from cell membrane and **(D)** nucleus. **(E)** Stream map with states identified for diverging branches S4 and S5 for the IgG1 anti PD-1 PC subsets. Volcano plot displayed for differential gene expression (p<0.05) across S4 and S5.

**Figure S4.**
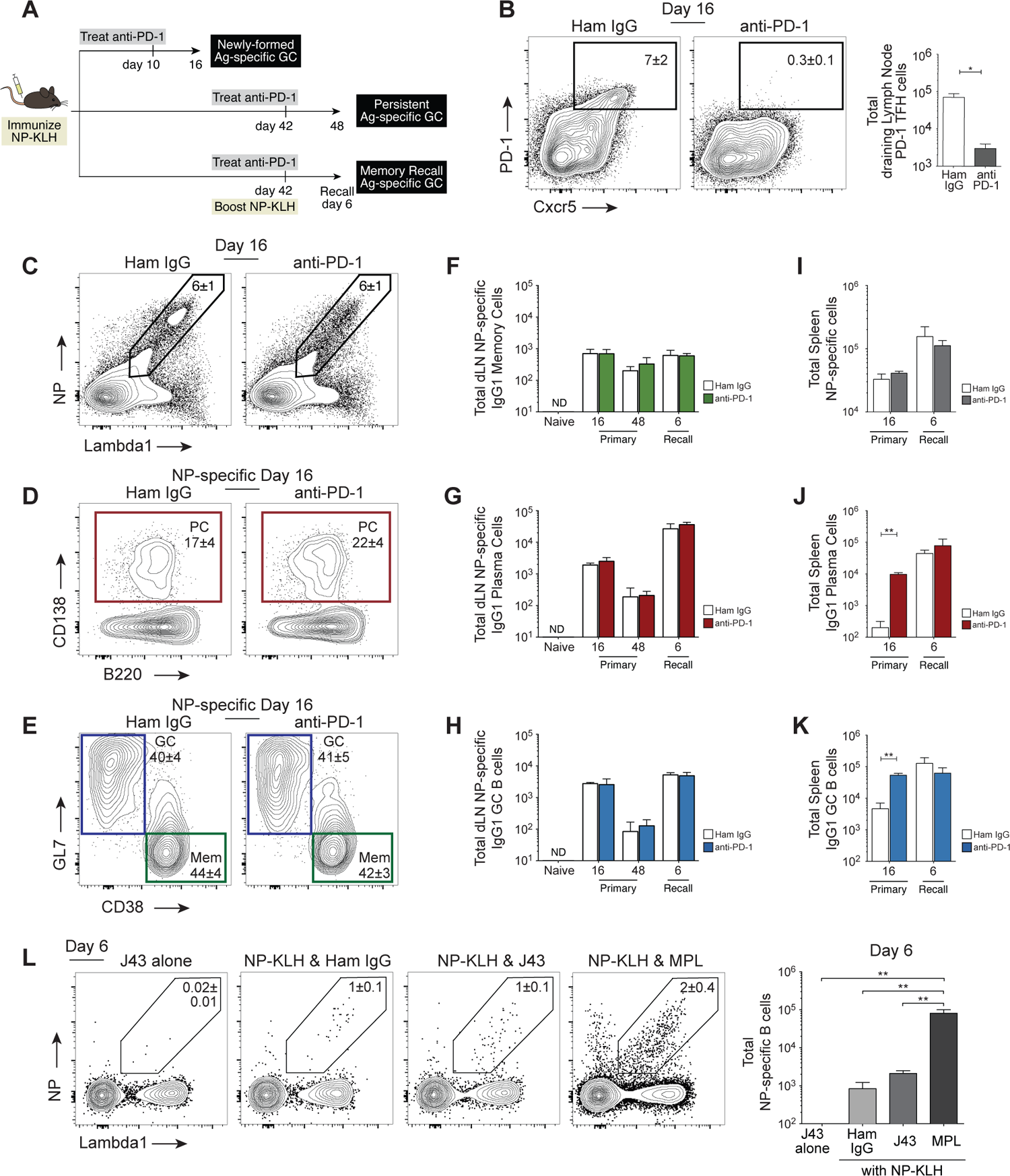
Vaccine-induced IgG1 GC B cells are not susceptible to PD-1 blockade. **(A)** Schematic of experimental procedure. Mice were immunized with NP-KLH in MPL adjuvant base of tail and treated with anti-PD-1 or isotype control antibody at the indicated timepoints. **(B)** Flow cytometry of activated CD4+ T cells (TCRb+ CD8-CD4+ CD44hi) in the draining lymph nodes following immunization and subsequent control IgG or anti-PD-1 treatment at day 10 with analysis at day 16. **(C)** Antigen-specific B cells (NP+ Lambda1+) from draining lymph nodes of (B), **(D)** antigen-specific PC, and **(E)** GC and memory B cells. **(F)** Quantification for the primary and recall response of total antigen-specific IgG1 memory B cell, **(G)** IgG1 PC, and **(H)** IgG1 GC B cells in the draining lymph nodes. **(I)** Total antigen-specific cells in the spleen. **(J)** All splenic IgG1 PC and **(K)** GC after control IgG or anti-PD-1 treatment. **(L)** Mice were immunized IP with anti-PD-1 (J43) alone, NP-KLH with isotype control antibody, NP-KLH with anti-PD1 (J43), or the NP-KLH with vaccine adjuvant TLR4 agonist (MPL). Antigen-specific B cells in the spleen with quantification (n=3) mean±SEM.

**Figure S5.**
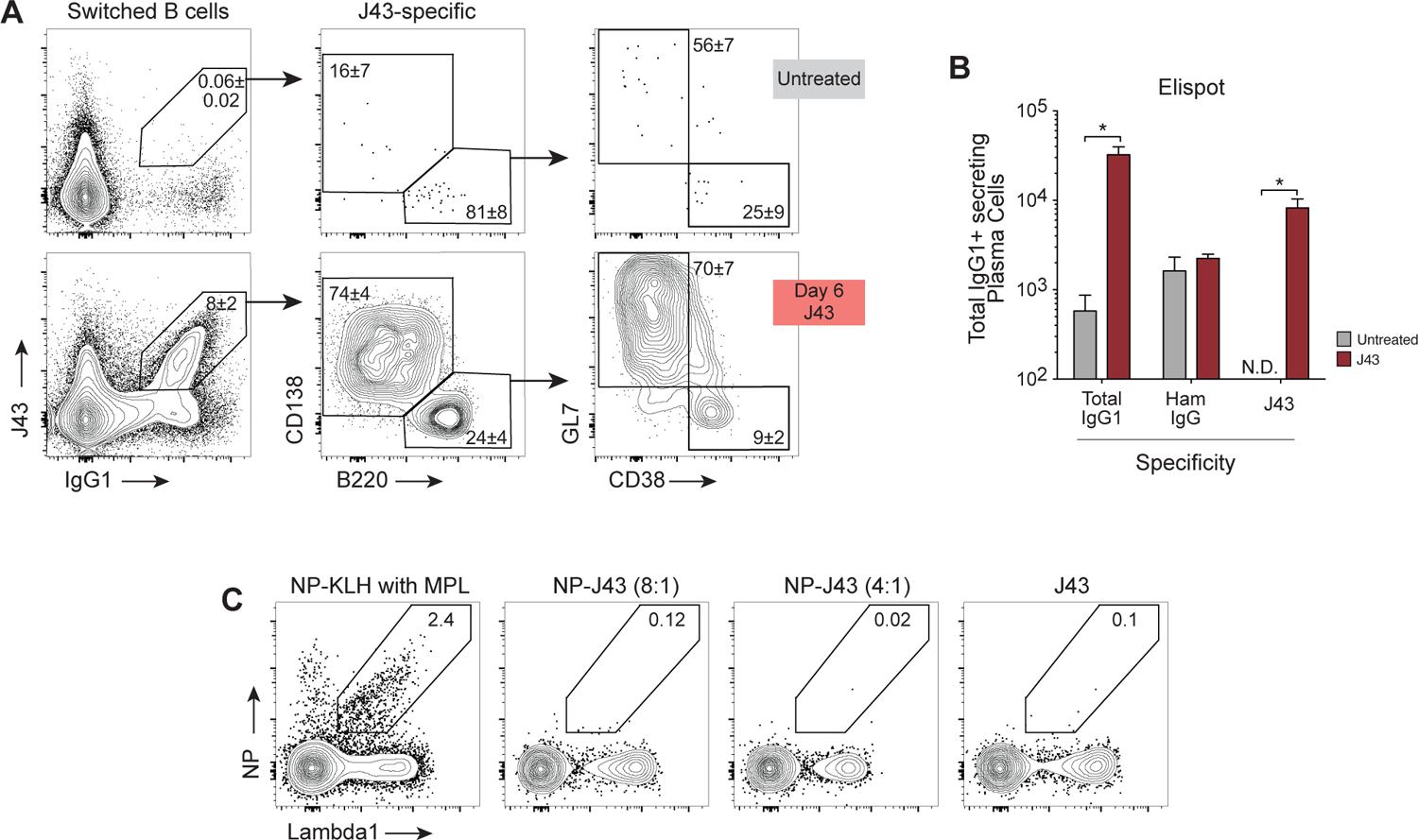
Pre-existing J43-specific IgG1 binding is enhanced with PD-1 blockade. **(A)** Steady state mice untreated or treated with anti-PD1 antibody (J43) and analyzed 6 days later for J43-binding. **(B)** Elispot quantification of specific binding to total IgG1, Ham IgG and J43 by secreting splenic B cells. **(C)** Mice treated with NP-KLH with MPL, NP-J43 (8:1 and 4:1 conjugation ratio) and J43 alone and assayed for antigen-specific B cells in the spleen.

